# Mitochondrial NAD+ import gates neuronal resilience to active SARM1

**DOI:** 10.64898/2026.10.05.756890

**Authors:** Lilianne Barbar, Anushka Bhat, Si Jia Chen, Amy Strickland, Sarah V. Schurr, Shea Sundali, Hayoung Kim, Yo Sasaki, A. Joseph Bloom, Jeffrey Milbrandt, Aaron DiAntonio

**Author notes:** Correspondence to Aaron DiAntonio.

## Abstract

SARM1 is an inducible NADase that executes pathological axon degeneration, but how NAD+ destruction causes neuronal failure remains unclear. Because SARM1 is predicted to act outside the mitochondrial matrix, cytosolic NAD+ loss has been viewed as the critical lethal lesion. We show that SARM1 activation rapidly depletes cytosolic and mitochondrial NAD+ and collapses ATP in both compartments, revealing early, coupled metabolic failure. Enhancing mitochondrial NAD+ import through SLC25A51, the mitochondrial NAD+ transporter, preserves mitochondrial NAD+, maintains mitochondrial function, and delays axon degeneration and neuronal death despite ongoing SARM1 activity. Unexpectedly, SLC25A51 restores cytosolic NAD+-dependent metabolism, reversing a glycolytic block and increasing NAD+ synthesis during stress. Glycolytic inhibition attenuates protection, identifying cytosolic metabolic recovery as an important component of neuronal resilience. AAV-SLC25A51 improves function in a chronic SARM1-dependent neuropathy model. Thus, mitochondrial NAD+ import gates SARM1-driven degeneration by limiting coupled mitochondrial-cytosolic metabolic failure, presenting a downstream intervention point for axon preservation.

## Introduction

SARM1 is an inducible NADase^1^ that is the central executioner of pathological axon degeneration^2,3^ and an important driver of neuronal cell death^4^. SARM1 is causally implicated in peripheral and optic neuropathies^5,6^, traumatic and toxic axon injury^7,8^, and CNS neurodegenerative disease^9,10^. Recent work has clarified how NAD+ metabolism regulates SARM1 activation: NAD+ opposes NMN-dependent activation via competitive binding to an allosteric site, with an increased NMN/NAD ratio favoring activation^11,12^. Once SARM1 is activated, NAD+ destruction leads to axon degeneration^1,13,14^. However, how NAD+ destruction results in irreversible neuronal failure remains unresolved, with recent studies identifying candidate downstream degenerative pathways^15–18^. SARM1 enzymatic activity is a druggable mechanism whose inhibition can block axon loss^19–21^. Therefore, defining the downstream metabolic lesions that makes SARM1 activation lethal is essential for understanding the determinants of neuronal resilience and how neuronal viability can be maintained despite SARM1 activation.

Cellular NAD+ is partitioned into nuclear, cytosolic, and mitochondrial pools. While NAD+ diffuses between the nucleus and cytosol through nuclear pores, mitochondrial matrix NAD+ is maintained as a distinct pool that, in neurons, accounts for a substantial fraction of total NAD+^22–25^. This compartmentalization is important when considering SARM1-mediated degeneration because current evidence places SARM1 outside the mitochondrial matrix.

SARM1 is mitochondria-associated, with evidence supporting localization at the outer mitochondrial membrane and more recent work suggesting that a portion resides in the intermembrane space and inner mitochondrial membrane^3,5^. In either case, SARM1 would be expected to consume cytosolic and/or intermembrane-space NAD+ rather than matrix NAD+. Thus, although current models emphasize cytosolic NAD+ depletion as the proximal metabolic lesion, it remains unresolved whether mitochondrial NAD+ is spared or secondarily depleted following SARM1 activation. It is also unknown whether mitochondrial NAD+ availability itself determines neuronal resilience to SARM1 activation.

Here we show that SARM1 activation rapidly depletes both cytosolic and mitochondrial NAD+, accompanied by ATP collapse in both compartments. The recent landmark discovery of the mammalian mitochondrial NAD+ importer SLC25A51^26–28^ allowed us to test whether mitochondrial NAD+ loss contributes functionally to degeneration. Overexpressing SLC25A51 increased and preserved mitochondrial NAD+ levels and delayed axon degeneration and neuronal death despite persistent SARM1 activity, demonstrating that SARM1 activation can be uncoupled from downstream degeneration. Unexpectedly, SLC25A51 also restored cytosolic NAD+-dependent metabolism. This was surprising because enhanced mitochondrial NAD+ import during ongoing SARM1-mediated NAD+ consumption might be expected to further limit cytosolic NAD+ availability. Instead, SLC25A51-mediated preservation of mitochondrial NAD+ sustained ATP levels and promoted stress-induced NAD+ biosynthesis, providing a route to maintain cytosolic NAD+-dependent reactions despite continued NADase activity. This protected state required glycolytic flux but did not require the mitochondrial NAD+ synthetase NMNAT3^29–31^, supporting a model in which imported mitochondrial NAD+ preserves broad cellular bioenergetics and enables compensatory extramitochondrial NAD+ synthesis. SLC25A51 overexpression also improved axonal physiology and motor function in an in vivo model of chronic SARM1-dependent neuropathy. Together, these findings demonstrate that SARM1 triggers degeneration via a coupled energetic, biosynthetic, and glycolytic failure gated by mitochondrial matrix NAD+ availability, and reveal how neuronal survival can be preserved despite ongoing SARM1 activity.

## Results

### SARM1 activation rapidly collapses cytosolic and mitochondrial NAD+ pools in neurons

Because activated SARM1 is thought to consume NAD+ outside the mitochondrial matrix, we directly compared cytosolic and mitochondrial NAD+ depletion following SARM1 activation. To establish a SARM1-dependent *in vitro* paradigm, we treated wild-type and *Sarm1* knockout cortical neurons with vacor^32^, a pharmacologic SARM1 activator, and quantified metabolites by LC-MS. In wild-type neurons, vacor induced accumulation of the SARM1 product cADPR^33^ and led to marked depletion of whole-cell NAD+ and ATP (p<.0001), whereas these responses were absent in *Sarm1* knockout neurons, confirming that the metabolic effects were SARM1-dependent (Fig. 1a). We then isolated cytosolic and mitochondrial fractions, which were confirmed to be enriched in HSP90 and COX4, respectively (Extended Data Fig. 1a), in order to quantify NAD+ in each compartment. Three hours of vacor treatment resulted in NAD+ depletion from both the cytosolic and mitochondrial fractions (p<.0001), indicating that mitochondrial NAD+ is lost early after SARM1 activation (Fig. 1b). Other metabolites measured in the same fractions, including cytosolic and mitochondrial FAD, inosine, and aspartate, were unchanged at this time (Extended Data Fig. 1b).

**Figure 1:**
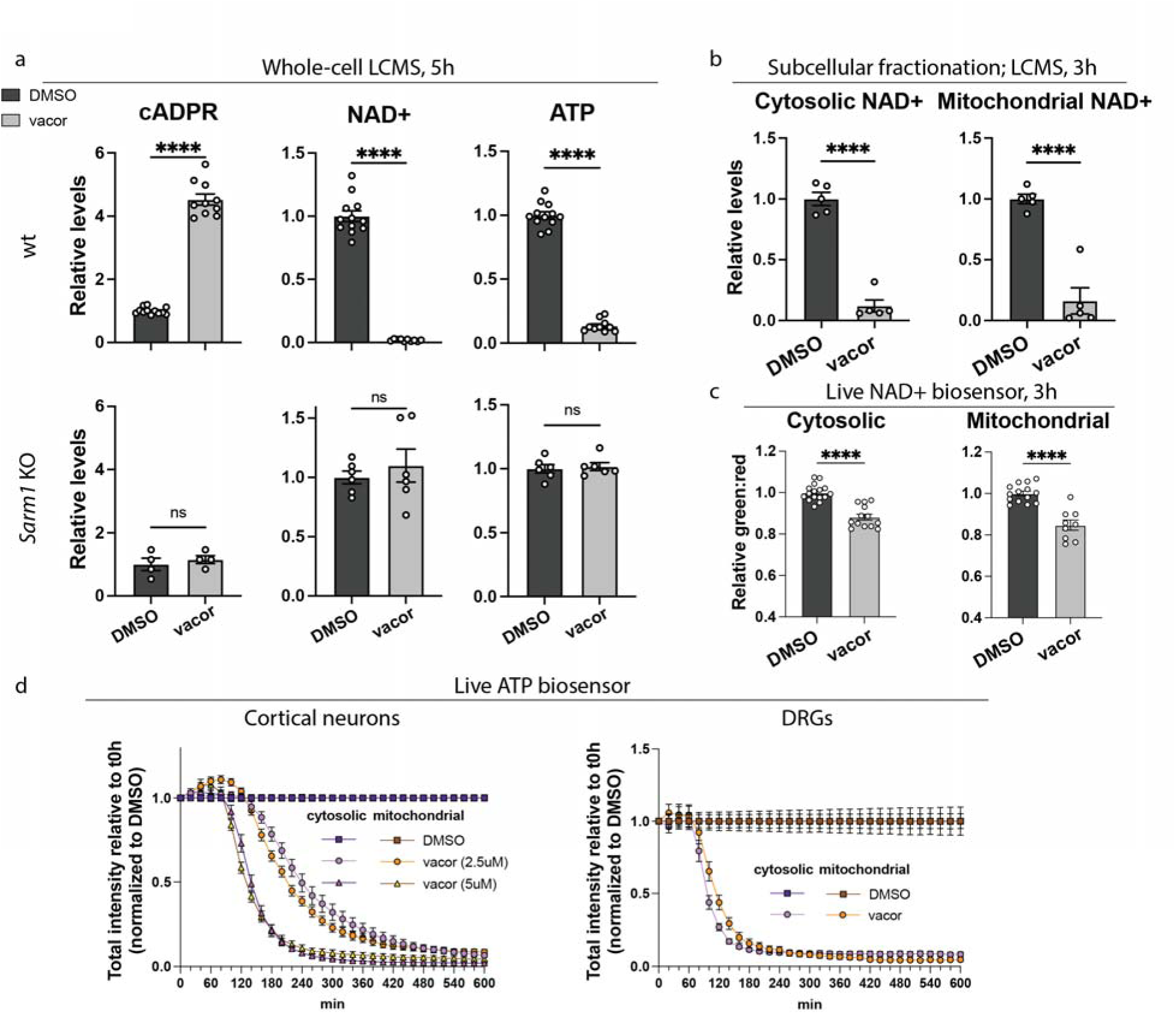
SARM1 activation rapidly collapses compartmental NAD+ and ATP in primary neurons. a. cADPR (cyclic adenosine diphosphate ribose), NAD+ (nicotinamide adenine dinucleotide), and ATP (adenosine triphosphate) levels from wild-type or *Sarm1*-KO primary mouse cortical neurons 5 hours following treatment with SARM1 activator vacor (5 µM). Metabolite levels are shown relative to DMSO-treated control, as measured by LC-MS. Error bars show mean ± SEM, with data points representing independently treated culture wells (wt: n=10-12; *Sarm1* KO: n=4-6) pooled across 3 independent culture preparations. Statistical significance was determined by two-tailed unpaired t-test (wt: p<.0001; SKO: p=.54 for cADPR, p=.52 for NAD+, p=.71 for ATP) b. Cytosolic and mitochondrial NAD+ levels in wild-type primary mouse cortical neurons treated with vacor (5 µM) vs DMSO for 3 hours. Metabolites are shown relative to DMSO-treated control, as measured by LC-MS following subcellular fractionation. Error bars show mean ± SEM, with data points representing independently treated culture wells (n=5) pooled across 3 independent culture preparations. Statistical significance was determined with a two-tailed unpaired t-test. c. Cytosolic and mitochondrial measurements of live NAD+ sensor FiNad in primary cortical neurons at 3 hours following 2.5 µM vacor vs DMSO treatment. Green:red ratio was measured with flow cytometry, and is shown relative to the ratio in DMSO-treated control. Increased green:red ratio reflects increased NAD+. Error bars show mean ± SEM, with data points representing independently treated culture wells (DMSO:n=14-16; vacor:n=9-13) pooled across 3 independent culture preparations. Statistical significance was determined with a two-tailed unpaired t-test. d. Live cytosolic and mitochondrial ATP microscopy imaging using the iATPSnFR2 sensor in primary cortical neurons (left) following treatment with vacor (2.5 µM; 5 µM) and in DRGs (right) following treatment with 12.5 µM vacor. Images were taken every 20 minutes and total signal intensity was measured and normalized to t0h and DMSO. Error bars show mean of independently treated culture wells, pooled across 3 independent culture preparations, ± SEM, with n=17-29 per group for cortical neurons and n= 25 for DRGs.

As an orthogonal live-cell readout, we expressed cytosolic or mitochondrially targeted ratiometric FiNad NAD+ sensors, which report NAD+-responsive green fluorescence normalized to an mCherry reference to adjust for sensor expression levels (Fig. 1c, Extended Data Fig. 1c), and quantified reporter ratios by flow cytometry^34^. We validated their sensitivity using neurons treated with the NAD+ biosynthesis inhibitor FK866 or the NAD+ precursor nicotinamide riboside (Extended Data Fig. 1d). As expected, FK866 decreased the FiNad signal, whereas nicotinamide riboside increased it in both compartments. Vacor reduced both cytosolic and mitochondrial FiNad signals (p<.0001, Fig. 1c) before signs of overt axon degeneration (Extended Data Fig. 1e), consistent with the NAD+ changes measured by LC-MS. As previously reported^34^, the FiNad sensor operates within a limited dynamic range, so modest changes in normalized fluorescence correspond to the substantially larger changes in LC-MS-measured NAD+ levels performed in parallel cultures (Fig. 1b-c). Together, these findings show that SARM1 activation rapidly depletes both cytosolic and mitochondrial NAD+ pools.

Because energetic failure is a central consequence of SARM1-mediated NAD+ depletion^35^, we next asked whether ATP loss parallels NAD+ depletion across the cytosolic and mitochondrial compartments. We expressed genetically encoded ATP reporters targeted to the cytosol or mitochondrial matrix to monitor ATP levels after vacor treatment^36^. Vacor triggered rapid ATP loss in both compartments, with cytosolic and mitochondrial ATP declining with similar kinetics in cortical and dorsal root ganglia (DRG) neurons (Fig. 1d, Extended Data Fig. 1f). The time course of ATP loss was consistent with the dramatic loss of whole cell ATP at 5 hours post vacor, as measured by LC-MS (Fig. 1a). This ATP loss was detectable before overt axon degeneration, which first became apparent at about 6 hours (Extended Data Fig. 1e), and progressed to near-complete depletion over the subsequent 4–6 hours, depending on vacor dose. Thus, SARM1 activation is accompanied by early loss of both cytosolic and mitochondrial ATP, indicating rapid disruption of compartmental energetic homeostasis. These findings suggested that mitochondrial metabolic failure may be an early and functionally important consequence of SARM1 activation, prompting us to ask whether preserving mitochondrial NAD+ could sustain energetics and alter the downstream course of SARM1-mediated degeneration.

### Mitochondrial NAD+ preservation delays degeneration despite ongoing SARM1 activation

We next asked whether enhancing SLC25A51-mediated mitochondrial NAD transport could preserve mitochondrial NAD+ during SARM1 stress. We overexpressed SLC25A51, the mammalian mitochondrial NAD+ transporter^26–28,37^, in primary neurons and first confirmed that it increased mitochondrial NAD+ under basal conditions (p<.0001, Fig. 2a). We also confirmed the specificity of this effect using 1) a transport-dead variant SLC25A51-K91A^26^, which did not increase mitochondrial NAD+ (p=.31), and 2) Ndt1, a heterologous mitochondrial NAD+ transporter from yeast^38^, which elevated mitochondrial NAD+ levels (p=.0004, Fig. 2a, Extended Data Fig. 2a). We next asked whether this SLC25A51 overexpression altered mitochondrial dysfunction, oxidative stress, and NAD+ loss downstream of SARM1 activation. Loss of mitochondrial NAD+ would be expected to impair mitochondrial energetics. Indeed, SARM1 activation led to a loss of mitochondrial membrane potential (p=.0001), and this effect was blunted by SLC25A51 overexpression, which resembled DMSO control (p=.98, Fig. 2b). SARM1 activation also triggered ROS accumulation in neurons, and this was robustly blocked by both a SARM1 chemical inhibitor^20^ and by SLC25A51 overexpression (Extended Data Fig. 2b).

**Figure 2:**
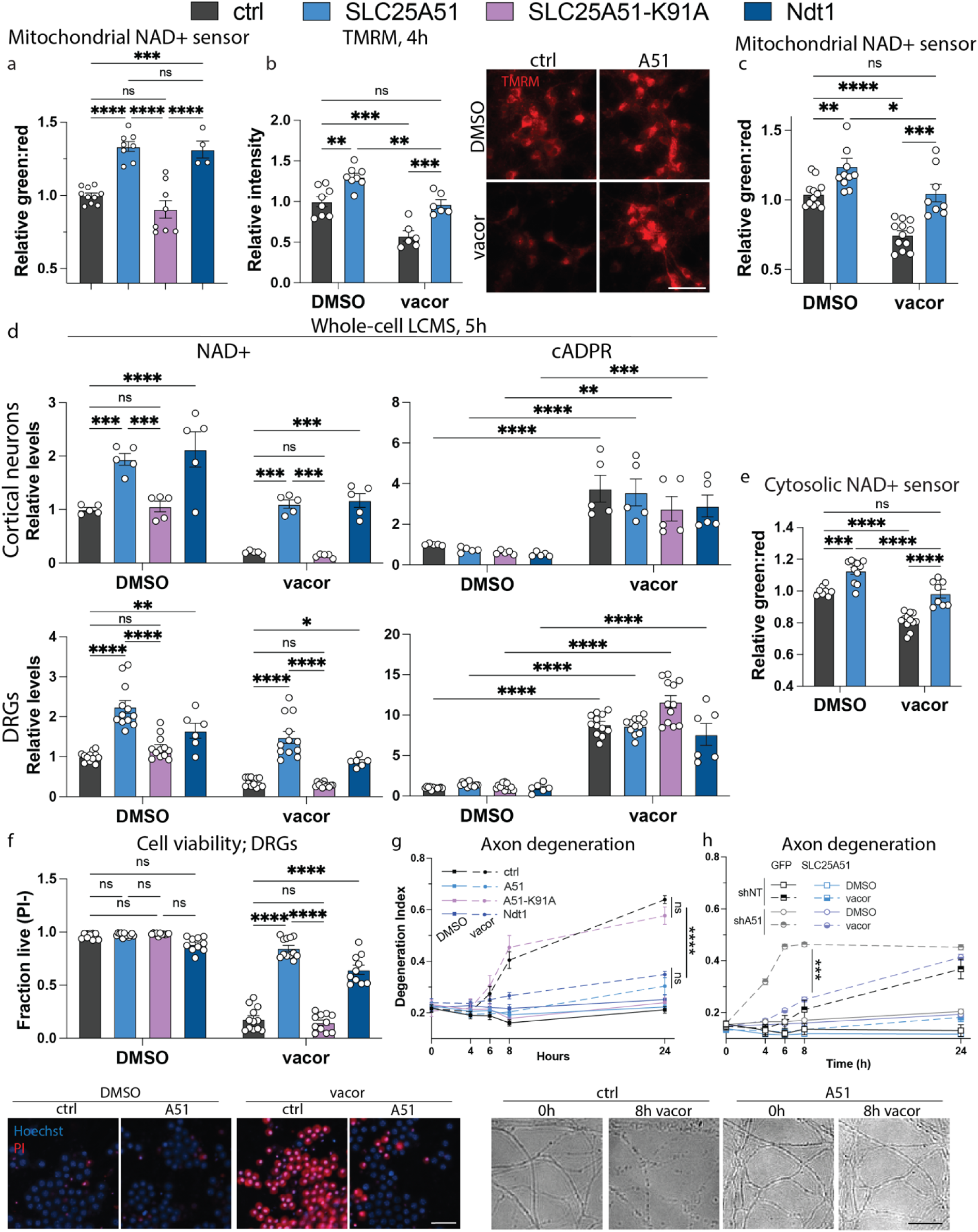
Mitochondrial NAD+ preservation with SLC25A51 lentiviral overexpression is neuroprotective downstream of SARM1 activation. a. Mitochondrial measurements of live NAD+ sensor FiNad in primary cortical neurons following lentiviral overexpression of wildtype SLC25A51 (A51) vs transport-dead variant SLC25A51-K91A (K91A) vs yeast mitochondrial transporter Ndt1 vs empty vector control. Green:red ratio was measured with flow cytometry, and is shown relative to the ratio in control cells transduced with empty vector. Error bars show mean ± SEM, with data points representing independently treated culture wells (n=10 for ctrl; 8 for A51 and K91A; 4 for Ndt1) pooled across 3 independent culture preparations. Statistical significance was determined by one-way ANOVA with Tukey’s multiple comparisons test. b. TMRM measurements in wild-type primary mouse cortical neurons transduced with A51 vs control (empty vector/GFP) and treated with vacor (2.5 µM) vs DMSO for 4 hours. Mean TMRM intensity was quantified from microscopy images and is shown relative to DMSO-treated control. Error bars show mean ± SEM, with data points representing independently treated culture wells (n=8 for DMSO; 6 for vacor) pooled across 3 independent culture preparations. Statistical significance was determined by 2-way ANOVA with Tukey’s multiple comparisons test. Representative images of TMRM (red) are shown at 4 hours following treatment with 2.5 µM vacor in control (empty vector) vs SLC25A51-overexpressing cortical neurons. Scale bar, 50 µm. c. Mitochondrial measurements of live NAD+ sensor FiNad in primary cortical neurons transduced with A51 vs empty vector control and treated with vacor (2.5 µM) vs DMSO. Green:red ratio was measured with flow cytometry, and is shown relative to the ratio in control cells transduced with empty vector. Error bars show mean ± SEM, with data points representing independently treated culture wells (n=8-12) pooled across 3 independent culture preparations. Statistical significance was determined by 2-way ANOVA with Tukey’s multiple comparisons test. d. NAD+ and cADPR levels from primary cortical neurons and DRG neurons transduced with A51, K91A, Ndt1, or control (empty vector/GFP), 5 hours following treatment with vacor (2.5 µM/12.5 µM). Metabolite levels are shown relative to DMSO-treated control, as measured by LC-MS. Error bars show mean ± SEM, with data points representing independently treated culture wells (cortical neurons: n=5; DRGs: n=12 for ctrl, SLC25A51, SLC25A51-K91A; n=6 for Ndt1) pooled across 3 independent culture preparations. Statistical significance was determined by 2-way ANOVA with Šídák’s multiple comparisons test. e. Cytosolic measurements of live NAD+ sensor FiNad in primary cortical neurons transduced with A51 vs empty vector control and treated with vacor (2.5 µM) vs DMSO. Green:red ratio was measured with flow cytometry, and is shown relative to the ratio in control cells transduced with empty vector. Error bars show mean ± SEM, with data points representing independently treated culture wells (n=8-12) pooled across 3 independent culture preparations. Statistical significance was determined by 2-way ANOVA with Tukey’s multiple comparisons test. f. Cell viability in primary DRG neurons transduced with A51, K91A, Ndt1, or GFP/empty vector control and treated with vacor (12.5 µM) for 24 hours. Representative images show propidium iodide (PI) in red and Hoechst (nuclei) in blue. Scale bar, 50 µm. Error bars show mean ± SEM, with data points representing independently treated culture wells (n=10-12) pooled across 3 independent culture preparations. Statistical significance was determined with a 2-way ANOVA with Tukey’s multiple comparisons test. g. Axon degeneration with representative images in primary DRG neurons transduced with A51 vs K91A vs Ndt1 vs control (empty vector or GFP) ± vacor (12.5 µM) vs DMSO. Degeneration index was quantified, with an increase indicating increased degeneration. Error bars show mean of independently treated culture wells, pooled across 3 independent culture preparations, ± SEM, with n=8-15. Statistical significance was determined by a mixed-effects model with Šídák’s multiple comparisons test, with pairwise comparisons shown for 24h. Scale bar, 25 µm. h. Axon degeneration in primary DRG neurons transduced with shRNA-resistant A51 or control (empty vector), in addition to non-targeting shRNA (shNT) or *Slc25a51-*targeting shRNA (shA51), and treated with vacor (12.5 µM) vs DMSO over 24 hours. Degeneration index was quantified, with an increase indicating increased degeneration. Error bars show mean of independently treated culture wells ± SEM, with n=5. Statistical significance was determined by a 2-way ANOVA with Tukey’s multiple comparisons test.

Furthermore, after vacor treatment, SLC25A51-expressing neurons maintained mitochondrial and whole-cell NAD+ levels at those of DMSO control neurons, avoiding the severe depletion otherwise induced by vacor (Fig. 2c-d). This protection did not reflect inhibition of SARM1 itself, as vacor still induced accumulation of the SARM1 enzymatic product cADPR in SLC25A51-expressing cortical and DRG neurons (Fig. 2d). Thus, SLC25A51 overexpression established a condition in which mitochondrial NAD+ was maintained at near basal levels of DMSO controls (p=.9995) despite ongoing SARM1 NADase activity at 5 hours post-vacor. Surprisingly, SLC25A51 also preserved cytosolic FiNad signal after vacor treatment (p=.96, Fig. 2e), indicating that SLC25A51 rescues both cytosolic and mitochondrial NAD+ levels. This was unexpected, given that increased mitochondrial NAD+ import might be predicted to deplete cytosolic NAD+ stores rather than restore them, a point which we explore further below.

We next asked if SLC25A51-mediated mitochondrial NAD+ preservation could protect neurons and axons despite continued SARM1 activity. SLC25A51 substantially delayed SARM1-dependent cortical neuron cell death (Extended Data Fig. 2c) and DRG neuron cell death and axon degeneration (Fig. 2f-g, Extended Data Fig. 2d), demonstrating that mitochondrial NAD+ preservation can uncouple persistent SARM1 activity from neuronal degeneration. To test whether the protection afforded by SLC25A51 was due to its mitochondrial NAD+ transport function, we also tested the effects of SLC25A51-K91A and Ndt1 in DRG neurons. After SARM1 activation by vacor, Ndt1 also preserved whole-cell NAD+, axon integrity, and neuronal viability similar to wild-type SLC25A51, whereas SLC25A51-K91A failed to prevent both metabolic and degenerative phenotypes (Fig. 2d, 2f-g). Thus, enhancing SLC25A51-mediated mitochondrial NAD+ transport is sufficient to preserve mitochondrial function and robustly delay degeneration downstream of persistent SARM1 activity. This protection depends upon NAD+ import rather than a transport-independent consequence of SLC25A51 overexpression.

Finally, we investigated whether loss of SLC25A51 would enhance SARM1 degenerative phenotypes. shRNA-mediated *Slc25a51* knockdown reduced Slc25a51 transcript levels by ∼90% (Extended Data Fig. 2e) and did not trigger spontaneous cell death or axon loss, but did accelerate axon degeneration and cell death in DRG neurons after vacor-induced SARM1 activation. This exacerbation caused by loss of endogenous SLC25A51 was rescued with lentiviral expression of an shRNA-resistant SLC25A51 construct (Fig. 2h, Extended Data Fig. 2f-g). SLC25A51 therefore serves as a neuronal resilience factor that supports neuron survival and axon integrity downstream of SARM1.

### Preservation of mitochondrial NAD+ reverses the broad metabolic consequences of SARM1 activation

SLC25A51 delayed degeneration despite persistent SARM1 activity, raising questions regarding how mitochondrial NAD+ preservation reshapes the downstream metabolic response to SARM1 activation. Using the compartment-targeted ATP reporters^36^, we found that SLC25A51 overexpression protects both cytosolic and mitochondrial ATP signals during vacor treatment, relative to vacor-treated control (p<.0001, Fig. 3a). Thus, SLC25A51-mediated mitochondrial NAD+ preservation has broader consequences, stabilizing both cytosolic and mitochondrial energetic homeostasis. LC-MS measurements similarly showed full protection against whole-cell ATP loss in both vacor-treated cortical and DRG neurons (Extended Data Fig. 3a). This raised the broader question of which metabolic pathways are preserved when mitochondrial NAD+ is maintained during ongoing SARM1-mediated NAD+ consumption.

**Figure 3:**
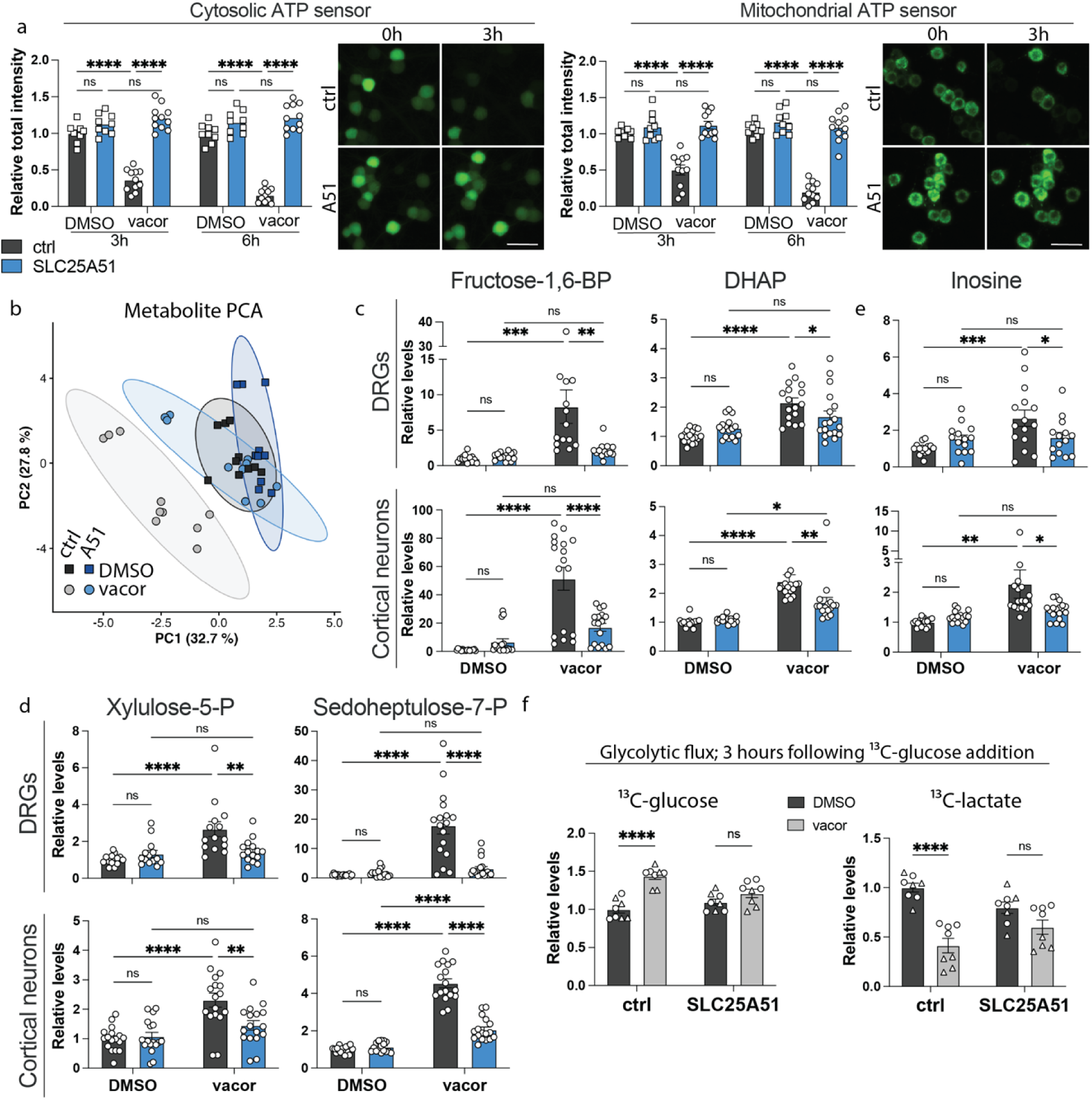
SLC25A51 restores global metabolic function downstream of SARM1 activation in primary neurons. a. Live cytosolic and mitochondrial ATP microscopy imaging using the iATPSnFR2 sensor in primary DRG neurons following treatment with vacor (12.5 µM) vs DMSO for 6 hours. Total signal intensity was measured and normalized to the DMSO-treated control. Error bars show mean ± SEM, with datapoints representing independently treated culture wells (n=9-11) pooled across 3 independent culture preparations. Statistical significance was determined by 2-way ANOVA with Šídák’s multiple comparisons test. Representative images of live cytosolic and mitochondrial ATP sensor in DRGs are shown at 0h and 3h following treatment with 12.5 µM vacor in control (empty vector) vs SLC25A51-overexpressing DRG neurons. Scale bar, 50 µm. b. Principal component analysis (PCA) based on 63 metabolomic features, demonstrating group separation across 4 conditions. Data points represent independently treated culture wells (n=10) pooled across 3 independent culture preparations. The shaded areas or ellipses represent the 95% confidence intervals. The percentages on the axes indicate the proportion of variance explained by each component. c. Levels of glycolytic intermediates fructose-1,6-bisphosphate and dihydroxyacetone phosphate (DHAP) in primary mouse dorsal root ganglia (DRG) and cortical neurons transduced with A51 vs empty vector control, 5 hours following treatment with vacor (12.5 µM for DRGs;2.5 µM for cortical neurons). Metabolite levels are shown relative to DMSO-treated control, as measured by LC-MS. Error bars show mean ± SEM, with data points representing independently treated culture wells (n=15-17) pooled across 3 independent culture preparations. Statistical significance was determined by 2-way ANOVA with Holm-Šídák’s multiple comparisons test. d. Levels of pentose phosphate pathway intermediates xylulose-5-phosphate and sedoheptulose-7-phosphate levels in primary mouse dorsal root ganglia (DRG) neurons and cortical neurons transduced with A51 vs empty vector control, 5 hours following treatment with vacor (12.5 µM for DRGs;2.5 µM for cortical neurons). Metabolite levels are shown relative to DMSO-treated control, as measured by LC-MS. Error bars show mean ± SEM, with data points representing independently treated culture wells (n=15-17) pooled across 3 independent culture preparations. Statistical significance was determined by 2-way ANOVA with Holm-Šídák’s multiple comparisons test. e. Purine metabolite inosine in primary DRG neurons and/or cortical neurons transduced with A51 vs control, 5 hours following treatment with vacor (12.5 µM for DRGs;2.5 µM for cortical neurons). Metabolite levels are shown relative to DMSO-treated control, as measured by LC-MS. Error bars show mean ± SEM, with data points representing independently treated culture wells (n=15-17) pooled across 3 independent culture preparations. Statistical significance was determined by 2-way ANOVA with Holm-Šídák’s multiple comparisons test. f. Relative levels of ^13^C-lactate and ^13^C-glucose in cortical neurons after incubating the cells with 1 mM ^13^C-glucose for 3 hours, starting 5 hours following treatment with vacor (2.5 µM) vs DMSO, as measured by LC-MS. Error bars show mean ± SEM, with data points representing independently treated culture wells (n=8) pooled across 2 independent culture preparations, which are represented by different datapoint symbols. Statistical significance was determined by a 2-way ANOVA with Tukey’s multiple comparisons test; Overexpression x drug treatment interaction was significant for both metabolites (p-values in main text).

To define broader metabolic consequences of SARM1 activation, we performed targeted metabolomic profiling of vacor-treated neurons, measuring 218 metabolites spanning glycolysis, the pentose phosphate pathway, the TCA cycle, and purine metabolism; 63 were consistently detected across neuronal samples and met quality-control criteria. Principal component analysis of these 63 reliably measurable metabolites showed that vacor-treated SLC25A51-overexpressing DRG neurons clustered with DMSO-treated control neurons (PERMANOVA; p=.615) and were distinct from vacor-treated control neurons (PERMANOVA; p=.008), indicating broad normalization of the vacor-induced metabolic state by SLC25A51 overexpression (Fig. 3b).

As expected, vacor treatment altered metabolites linked to mitochondrial and cytosolic NAD+-dependent metabolism. Among mitochondrial metabolites, malate accumulated markedly after SARM1 activation, consistent with impaired flux through malate dehydrogenase, which requires NAD+ to oxidize malate to oxaloacetate (Extended Data Fig. 3b-c). Vacor also induced a modest increase in alpha-ketoglutarate (Extended Data Fig. 3b-c). SLC25A51 overexpression prevented both abnormalities. These metabolite changes, along with the TMRM and mitochondrial ATP sensor data, demonstrate that SARM1 activation produces substantial mitochondrial metabolic abnormalities that are counteracted by SLC25A51 overexpression.

Among the most prominent vacor-induced pathway-level changes was a disruption of glycolysis. Metabolites upstream of the NAD+-dependent glyceraldehyde-3-phosphate dehydrogenase step accumulated, consistent with a pronounced glycolytic bottleneck developing after SARM1 activation (Fig. 3c, Extended Data Fig. 3d-e). This pattern could explain how NAD+ depletion propagates into broader energetic failure. Consistent with the unexpected rescue of cytosolic FiNad signal during SARM1 activity (Fig. 2e), SLC25A51 overexpression substantially corrected this vacor-induced glycolytic bottleneck in both DRG and cortical neurons, normalizing the levels of metabolites upstream of this NAD+-dependent glycolytic step including Fructose-1,6-BP, DHAP, Glucose-6-P, and Fructose-6-P (Fig. 3c, Extended Data Fig. 3e).

Because glycolytic intermediates downstream of GAPDH were difficult to reliably quantify, we independently assayed glycolytic flux with ^13^C-labelled glucose. In control cortical neurons, SARM1 activation by vacor caused intracellular ^13^C-glucose to accumulate, while the ^13^C-lactate labeling was reduced by ∼60%. This pattern is consistent with glycolytic inhibition, as ^13^C-glucose accumulates rather than being converted to ^13^C-lactate. SLC25A51 overexpression attenuated both changes, with a significant SLC25A51 overexpression x treatment interaction for each metabolite (^13^C-lactate, p =.0045; ^13^C-glucose, p =.0019; two-way ANOVA) (Fig. 3f). The lactate:pyruvate ratio was unchanged by vacor, arguing against a change in LDH equilibrium as the basis for the reduced 13C3-lactate signal (Extended Data Fig. 3h).

In addition to glycolysis and mitochondrial metabolism, vacor altered metabolites in several interconnected cytosolic pathways. Intermediates in the non-oxidative pentose phosphate pathway and purine metabolism were strongly perturbed, including a large increase in xylulose-5-phosphate and sedoheptulose-7-phosphate and marked accumulation of inosine, succinyladenosine, hypoxanthine, AMP, and IMP upon SARM1 activation (Fig. 3d-e, Extended Data Fig. 3f-g). SLC25A51 expression normalized these pentose phosphate and purine abnormalities in both DRG and cortical neurons (Fig. 3d-e, Extended Data Fig. 3f-g), indicating that cytosolic metabolic function can be improved despite persistent SARM1 catalytic activity. Hexosamine biosynthesis intermediate N-acetylglucosamine-1-phosphate also accumulated following vacor treatment, consistent with the GAPDH bottleneck observed, and this was prevented with SLC25A51 overexpression (Extended Data Fig. 3i). Finally, the rescue of cytosolic ATP levels with SLC25A51 overexpression in the presence of activated SARM1 raised the question of whether cytosolic ATP-dependent processes had been functionally restored. We found that ureidosuccinic acid is depleted after vacor-induced SARM1 activation (Extended Data Fig. 3j). The rate-limiting step in its synthesis is ATP-dependent. SLC25A51 overexpression rescued ureidosuccinic acid levels after vacor treatment, suggesting a restoration of the upstream cytosolic ATP-dependent reaction.

Together, these findings show that mitochondrial NAD+ preservation prevents broad SARM1-induced metabolic disruption across mitochondrial and cytosolic pathways in both DRG and cortical neurons. In particular, in addition to rescuing mitochondrial membrane potential and mitochondrial NAD+ and ATP loss, SLC25A51 restores both cytosolic NAD+-dependent glycolytic metabolism and cytosolic ATP-dependent reactions, supporting a model in which SLC25A51-mediated preservation of mitochondrial NAD+ stabilizes broad cellular metabolic homeostasis and thereby promotes neuronal resilience downstream of SARM1 activation.

### Mitochondrial NAD+ preservation promotes NAD+ resynthesis and reveals a requirement for glycolytic metabolism

The rescue of cytosolic NAD+ and NAD+-dependent metabolism by SLC25A51-mediated mitochondrial NAD+ preservation raised two questions: how is cytosolic metabolic function restored, and does this restoration contribute to axon protection? Because NMNAT-dependent NAD+ synthesis requires ATP, we considered whether preservation of mitochondrial energetics and cytosolic ATP might support continued NAD+ synthesis despite ongoing SARM1-mediated NAD+ hydrolysis (Fig. 4a). To test this, we tracked NAD+ synthesis using D4-labeled nicotinamide (D4-NAM)^39^. After vacor treatment, SLC25A51 robustly increased D4-NAM incorporation into NAD+, producing approximately twice as much labeled NAD+ as control neurons (Fig. 4b) in both DRGs (p=.0004) and cortical neurons (p<.0001). Thus, preservation of mitochondrial NAD+ levels promotes NAD+ synthesis and replenishment under conditions of SARM1-mediated metabolic stress.

**Figure 4:**
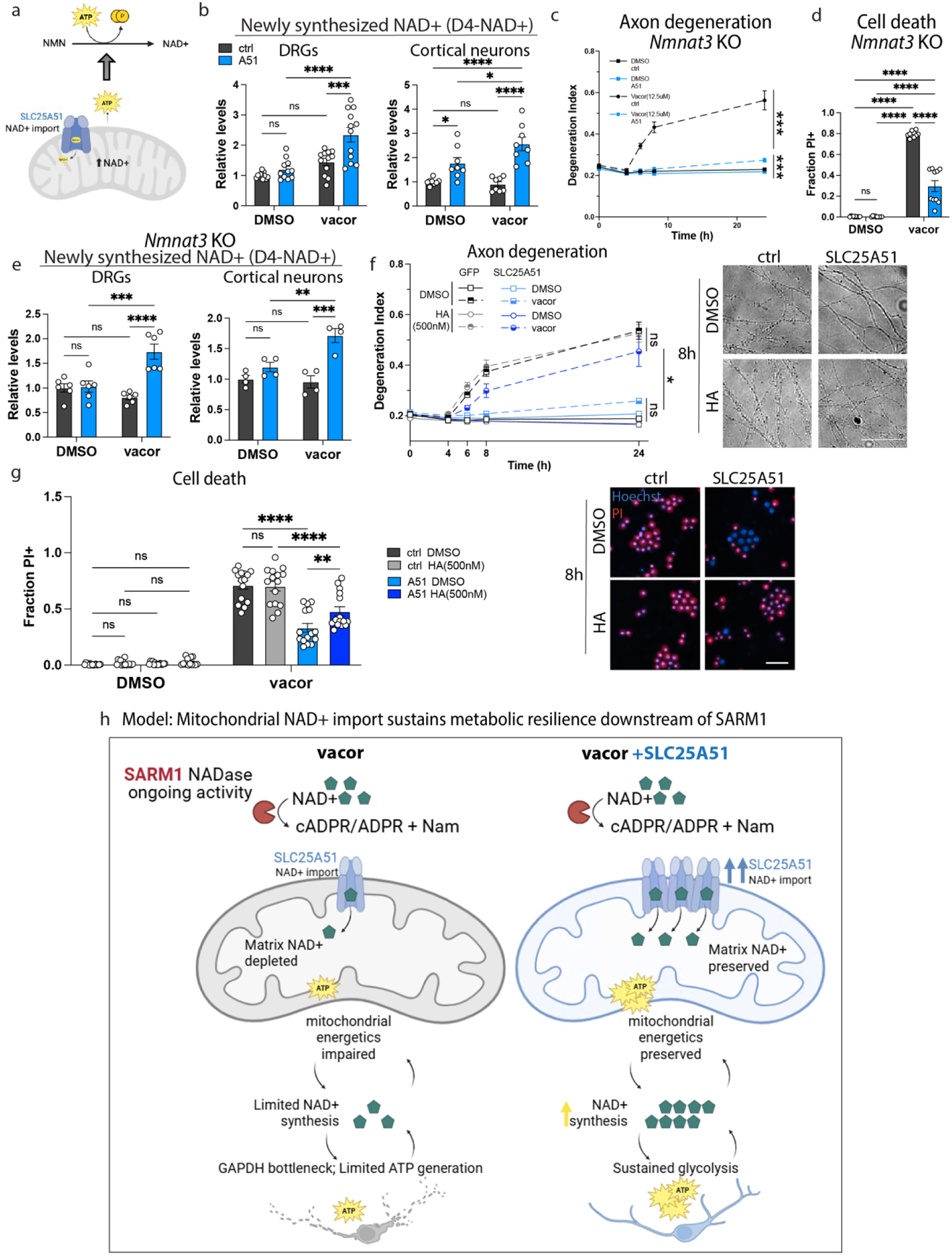
SLC25A51 boosts neuronal NAD+ synthesis downstream of SARM1, coupling cytosolic and mitochondrial metabolism to promote survival. a. Schematic illustrating the proposed mechanism by which mitochondrial NAD+ preservation by SLC25A51 may support cytosolic NAD+ resynthesis downstream of SARM1 activation, via ATP-dependent NMNAT, coupling mitochondrial and cytosolic NAD+ maintenance and metabolism. Created in BioRender. DiAntonio, A. (2026) https://BioRender.com/a5n43wu b. D4-labeled NAD+ levels in primary DRG neurons and cortical neurons transduced with A51 vs control, following treatment with vacor (12.5 µM/2.5 µM) vs DMSO for 2.5 hours in the presence of D4-nicotinamide (2,4,5,6-deuterium Nam; 300 µM). Labeled NAD+ levels are shown relative to DMSO-treated control, as measured by LC-MS. Error bars show mean ± SEM, with data points representing independently treated culture wells (n=11-12 for DRGs; n=8 for cortical neurons) pooled across 3 independent culture preparations. Statistical significance was determined by 2-way ANOVA with Tukey’s multiple comparisons test. c. Axon degeneration in primary *Nmnat3*-knockout DRG neurons transduced with A51 vs control, treated with vacor (12.5 µM) vs DMSO over 24 hours. Degeneration index was quantified, with an increase indicating increased degeneration. Error bars show mean of independently treated culture wells, pooled across 2 independent culture preparations, ± SEM, with n=10. Statistical significance was determined by a 2-way ANOVA with Tukey’s multiple comparisons test with pairwise comparisons shown for 24h. d. Cell death in primary *Nmnat3*-knockout DRG neurons transduced with A51 or control (GFP/empty vector) and treated with vacor (12.5 µM) for 24 hours, measured by the fraction of nuclei positive for propidium iodide (PI). Error bars show mean ± SEM, with data points representing independently treated culture wells (n=10) pooled across 2 independent culture preparations. Statistical significance was determined with a 2-way ANOVA with Tukey’s multiple comparisons test. e. D4-labeled NAD+ levels in *Nmnat3* KO primary DRG and cortical neurons transduced with A51 vs empty vector control, following treatment with vacor (12.5 µM) vs DMSO for 2.5 hours in the presence of D4-nicotinamide (2,4,5,6-deuterium Nam; 300 µM). Labeled NAD+ levels are shown relative to DMSO-treated control, as measured by LC-MS. Error bars show mean ± SEM, with data points representing independently treated culture wells (n=4-6) pooled across 2 independent culture preparations. Statistical significance was determined by 2-way ANOVA with Tukey’s multiple comparisons test. f. Axon degeneration with representative images in primary DRG neurons transduced with A51 vs control, treated with vacor (12.5 µM) ± glycolytic inhibitor heptelidic acid (HA; 500 nM) vs DMSO over 24 hours. Degeneration index was quantified, with an increase indicating increased degeneration. Error bars show mean of independently treated culture wells, pooled across 3 independent culture preparations, ± SEM, with n=14-15. Statistical significance was determined by 2-way ANOVA with Tukey’s multiple comparisons test with pairwise comparisons shown for 24h. Scale bar, 50 µm. g. Cell death with representative images in primary DRG neurons transduced with A51 or control (GFP/empty vector) and treated with vacor (12.5 µM) ± glycolytic inhibitor heptelidic acid (HA; 500 nM) vs DMSO for 24 hours, measured by the fraction of nuclei positive for propidium iodide (PI). Error bars show mean ± SEM, with data points representing independently treated culture wells (n=15) pooled across 3 independent culture preparations. Statistical significance was determined with a 2-way ANOVA with Tukey’s multiple comparisons test. Scale bar, 50 µm. h. Model for SLC25A51-mediated metabolic resilience downstream of SARM1. SARM1 activation consumes cytosolic NAD+ and produces coupled cytosolic and mitochondrial metabolic failure. Cytosolic NAD+ depletion is accompanied by mitochondrial NAD+ depletion, loss of mitochondrial and cytosolic ATP, impaired NAD+ replenishment, a glycolytic bottleneck, and neuronal degeneration. These changes reinforce each other, as NAD+ is required for glycolysis and oxidative phosphorylation, while ATP is required for NAD+ synthesis. SLC25A51 overexpression enhances mitochondrial NAD+ transport and preserves matrix NAD+ during ongoing SARM1 activity. Maintenance of mitochondrial function and ATP availability is proposed to support NMNAT-dependent NAD+ replenishment, which in turn sustains glycolysis and further ATP production. Together, these coupled mitochondrial and cytosolic metabolic effects delay SARM1-dependent axon degeneration and neuronal death. Created in BioRender. DiAntonio, A. (2026) https://BioRender.com/8tdx5su

Because this recovery could reflect mitochondrial or extramitochondrial NAD+ synthesis, we tested whether protection required NMNAT3, the mitochondrial NAD+ synthetase. NMNAT3 KO mice are viable with no obvious neurological phenotypes^40,41^ and *Nmnat3* knockout neurons are healthy *in vitro*. SLC25A51 remained protective against SARM1-dependent axon degeneration and cell death in *Nmnat3* knockout DRG neurons, demonstrating that mitochondrial NAD+ synthesis is dispensable for SLC25A51-mediated protection (Fig. 4c-d).

After vacor treatment, SLC25A51 also increased D4-NAM incorporation into NAD+ relative to control DRG (p<.0001) and cortical (p=.0003) neurons in the absence of NMNAT3, demonstrating that NMNAT3 is not required for the SLC25A51-mediated increase in newly synthesized NAD+ downstream of SARM1 (Fig. 4e). To test our hypothesis that extramitochondrial NAD+ synthesis is necessary for SLC25A51-mediated neuroprotection, we assessed the role of SLC25A51 overexpression in axotomy, where the highly labile cytosolic NMNAT2 is rapidly lost from the distal axon^42,43^. If SLC25A51-mediated protection requires extramitochondrial NAD+ synthesis, SLC25A51 should not protect from axotomy-induced axon degeneration since neither nuclear NMNAT1 nor the cytosolic NMNAT2 is available in the distal axon within a few hours of axotomy. Consistent with our prediction, SLC25A51 overexpression did not inhibit degeneration of distal axons following axotomy (Extended Data Fig. 4a), which supports the model that SLC25A51-mediated protection requires extramitochondrial NAD+ synthesis.

We next asked whether restored glycolytic metabolism contributes functionally to protection. If restored glycolytic flux is a component of the protected state rather than a byproduct of it, then blocking glycolysis should reduce the protection afforded by SLC25A51 overexpression. To do this, we identified a dose of the GAPDH inhibitor heptelidic acid that does not cause axon degeneration or enhance vacor-induced degeneration in control neurons. Nevertheless, this dose does strongly reduce the protection from vacor-induced axon degeneration and cell death afforded by SLC25A51 overexpression (Fig. 4f-g). Thus, SLC25A51-mediated rescue is partly dependent on preserving glycolytic flux downstream of SARM1 activation. Given that SLC25A51 also preserves mitochondrial ATP, we asked whether mitochondrial ATP production contributes more generally to neuronal survival following SARM1 activation. Low-dose oligomycin, an inhibitor of mitochondrial ATP synthase, exacerbated vacor-induced degeneration and reduced SLC25A51-mediated protection without triggering degeneration on its own (Extended Data Fig. 4b-c). Taken together with the preservation of ATP and increased accumulation of newly synthesized NAD+ described above, these findings support a model in which mitochondrial NAD+ preservation promotes cytosolic recovery by supporting NAD+ synthesis and glycolytic metabolism, allowing neurons to remain viable during ongoing SARM1 activation (Model, Fig. 4h).

### SLC25A51 preserves axonal function in a chronic SARM1-dependent neuropathy in vivo

To determine whether mitochondrial NAD+ preservation can modify neuronal dysfunction *in vivo* under conditions of chronic endogenous metabolic stress, we expressed SLC25A51 in the *Nmnat2*^V98M/R232Q^ neuropathy model, in which impaired axonal NMNAT2 activity produces a progressive, SARM1-dependent distal axon degeneration^44^. The degeneration in this model is driven by neuronal SARM1, as neuronal expression of a dominant-negative SARM1 rescues the neuropathy. Mice were injected with AAV intrathecally between postnatal days 13 and 19; AAV-mediated SLC25A51 delivery under the strong, constitutive EF1a promoter produced robust transgene expression in the spinal cord (Extended Data Fig. 5a). SLC25A51 expression improved motor performance until at least six months of age (p=.0002, treatment main effect by 2-way ANOVA), as measured by the inverted screen test (Fig. 5a). Electrophysiologic analysis similarly demonstrated improved compound muscle action potentials, consistent with preservation of functional motor axons (Fig. 5b), and tibialis anterior muscle mass was increased (p=.0444), indicating reduced denervation-associated atrophy (Fig. 5c). Expression of the repair Schwann cell marker c-Jun was reduced in SLC25A51-treated nerves (p=.036), consistent with reduced denervation-associated repair Schwann cell formation (Fig. 5d, Extended Data Fig. 5b). Finally, histologic analysis showed modest preservation of axon density in the sciatic nerve of SLC25A51-treated mice relative to control GFP-treated mice (p=.0183), suggesting that SLC25A51 may functionally support metabolically stressed axons rather than fully blocking axon degeneration (Fig. 5e). In this neuropathy model, levels of the SARM1 biomarker cADPR are elevated^44^, and these levels are unchanged by overexpression of SLC25A51 compared to the GFP control (p=.87, Fig. 5f). NAD+ levels in the nerve were modestly increased by SLC25A51 overexpression (Extended Data Fig. 5c). Hence, SLC25A51 is neuroprotective without suppressing SARM1 activation.

**Figure 5:**
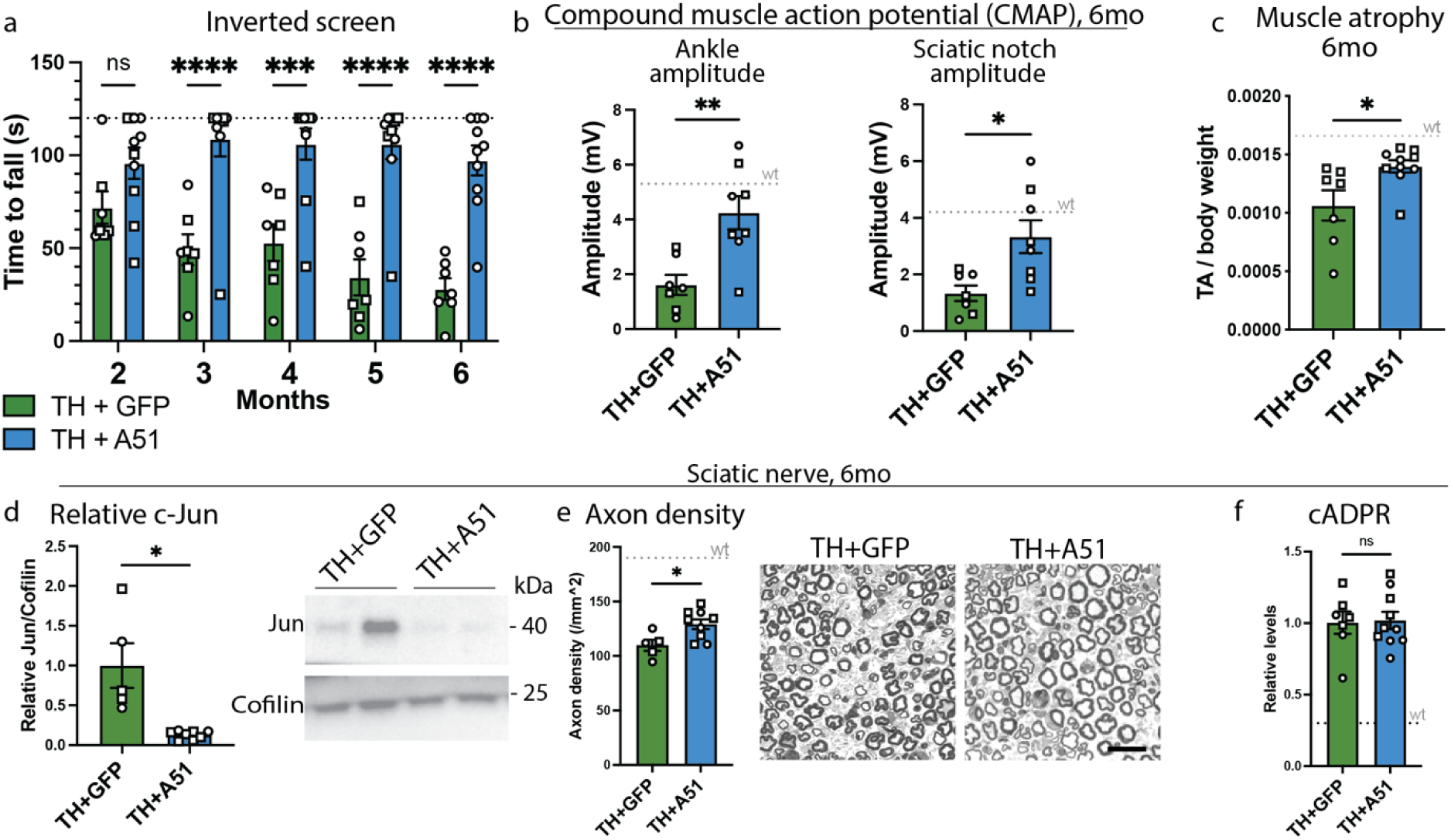
SLC25A51 promotes *in vivo* resilience and delays progression in a SARM1-dependent neuropathy mouse model. a. Motor performance of *Nmnat2*^V98M/R232Q^ (trans-het; TH) mice injected between postnatal days 13 and 19 with AAV9/PHP.eB/S dual capsid virus expressing SLC25A51 or GFP under the EF1α promoter, assessed by inverted screen testing at 2,3,4,5, and 6 months of age. Latency to fall is shown as mean ± SEM, with data points representing individual animals (n=7 for GFP; n=10 for A51). Statistical significance was determined by 2-way ANOVA with Šídák’s multiple comparisons test. Males are represented as squares and females as circles. b. Compound muscle action potential (CMAP) amplitude in TH mice injected with AAV-SLC25A51 vs AAV-GFP, measured at 6 months of age, when stimulating at the ankle or at the sciatic notch (SN). Error bars show mean ± SEM, with data points representing individual animals (n=7-8). Statistical significance was determined with two-tailed Welch’s t-test (p=.0032 for ankle stimulation; p=.0107 for sciatic notch stimulation). Males are represented as squares and females as circles. c. Tibialis anterior muscle mass in TH mice injected with AAV-SLC25A51 vs AAV-GFP, measured at 6 months of age. Muscle mass is shown normalized to body weight. Error bars show mean ± SEM, with data points representing individual animals (n=7 for GFP; n=10 for A51). Statistical significance was determined by two-tailed Welch’s t-test. Males are represented as squares and females as circles. d. Western blot confirmation of reduced levels of repair Schwann cell marker c-Jun in 6 month sciatic nerves of TH mice injected with AAV-SLC25A51 vs GFP control. Quantification of c-Jun band intensity was normalized to loading control Cofilin and GFP control. Error bars show mean ± SEM, with data points representing individual animals (n=5 for GFP; n=8 for A51). Statistical significance was determined with two-tailed Welch’s t-test. Males are represented as squares and females as circles. e. Representative histological images and quantification of axon density in sciatic nerve sections from TH mice expressing AAV-FLAG-SLC25A51 vs AAV-GFP at 6 months of age. Axon density was quantified with toluidine blue. Error bars show mean ± SEM, with data points representing individual animals (n=5 for GFP; n=9 for A51). Statistical significance was determined with two-tailed Welch’s t-test. Males are represented as squares and females as circles. Scale bar, 20 µm. f. cADPR levels in sciatic nerves of TH mice injected with AAV-SLC25A51 vs AAV-GFP, measured at 6 months of age. Metabolite levels are shown relative to GFP-treated control, as measured by LC-MS. Error bars show mean ± SEM, with data points representing individual animals (n=7 for GFP; n=10 for A51). Statistical significance was determined with two-tailed Welch’s t-test. Males are represented as squares and females as circles.

Because this model displays chronic axonal NAD+ insufficiency caused by reduced NMNAT2 function, the ability of SLC25A51 to improve motor performance, axonal physiology, muscle preservation, and injury-associated Schwann cell responses indicates that SLC25A51-mediated mitochondrial NAD+ preservation remains protective even when endogenous NAD+ synthetic capacity is limited and the upstream SARM1-dependent degenerative program remains engaged. These findings extend the *in vitro* observation that SLC25A51-mediated mitochondrial NAD+ preservation delays degeneration downstream of SARM1 and support mitochondrial NAD+ availability as a determinant of neuronal resilience under chronic NAD+ stress in vivo.

## Discussion

Our findings define a mechanism by which SARM1 activation promotes axon and neuron death by identifying mitochondrial NAD+ loss as an early and functionally important step in the downstream degenerative cascade. A substantial body of work implicates mitochondria in SARM1-dependent degeneration: mitochondrial dysfunction can promote SARM1 activation^33,45,46^, conversely SARM1 activation rapidly produces mitochondrial dysfunction^35,47^, and in chronic neuropathy these reciprocal effects can form a feed-forward loop that amplifies axonal pathology^6^. What remained unresolved was how SARM1-mediated NAD+ loss propagates into the metabolic failure that ultimately drives degeneration. SARM1-mediated degeneration has often been viewed as a consequence of global NAD+ depletion, with many NAD+-dependent reactions collapsing in parallel until axons can no longer maintain homeostasis^15–17,48^. Our data sharpen this model by showing that, while SARM1 rapidly depletes both mitochondrial and cytosolic NAD+, SLC25A51-mediated preservation of mitochondrial NAD+ potently delays degeneration despite persistent SARM1 activity. Thus, mitochondrial NAD+ availability gates the coupled energetic, biosynthetic, and glycolytic failure that drives neuronal degeneration.

Metabolomic profiling further supports this model by revealing disturbances extending beyond mitochondrial metabolism to glycolysis, the pentose phosphate pathway (PPP), and purine metabolism, which draws on PPP-derived ribose-5-phosphate. The accumulation of metabolites upstream of NAD+-dependent dehydrogenase reactions is consistent with impaired flux through multiple NAD+-dependent steps. Direct flux measurements confirmed impaired glycolysis, consistent with the GAPDH bottleneck indicated by the metabolomic profile. Some of the accumulated metabolites may also arise directly from SARM1 reaction-product disposal: SARM1 cleavage of NAD+ generates ADP-ribose, which is hydrolyzed to AMP and ribose-5-phosphate by Nudix hydrolases such as NUDT5^49^ and NUDT9^50^, linking NAD+ cleavage to both purine and PPP metabolism. From either source, the resulting accumulation of phosphorylated glycolytic and PPP intermediates, together with accumulating adenylates, could further exacerbate energy failure by sequestering inorganic phosphate and promoting ATP-consuming futile cycles^51,52^. Consistent with a pathogenic contribution from these disposal products, an unbiased forward genetic screen in Drosophila recovered loss of CG4098, the NUDT9 homolog, as a strong suppressor of neurodegeneration caused by loss of dNmnat^53^. Thus, the intermediates that accumulate as a result of the NAD+ bottlenecks following SARM1 activation may further constrain the ATP availability needed to relieve them. SLC25A51 overexpression prevents these metabolic changes broadly, identifying matrix NAD+ availability as a shared upstream constraint on these cytosolic pathways.

The mechanisms linking mitochondrial NAD+ import to cytosolic metabolic recovery likely reflect reciprocal coupling among ATP production, NAD+ synthesis, and glycolytic flux (see Model, Fig. 4h). SARM1-mediated NAD+ consumption directly limits cytosolic NAD+-dependent reactions, including the GAPDH step of glycolysis, and impairs oxidative phosphorylation by disrupting mitochondrial metabolism and membrane potential. These mechanisms drive SARM1-dependent ATP depletion. NMNAT-dependent NAD+ synthesis^54^ requires ATP, and so this creates a self-reinforcing failure state: NAD+ loss impairs glycolysis and oxidative metabolism, ATP loss limits NAD+ resynthesis, and declining NAD+ availability further constrains cytosolic metabolism. By preserving mitochondrial NAD+, SLC25A51 maintains mitochondrial function and ATP availability, allowing cytosolic NAD+ biosynthesis to continue despite ongoing SARM1 activity (see Model, Fig. 4h). Because NAMPT is inhibited by NAD+^55^, SARM1-mediated NAD+ depletion may also relieve feedback inhibition on the salvage pathway and so promote NAD+ resynthesis. This may act alongside preserved ATP availability to enable NAD+ resynthesis, although we did not directly test this mechanism. This ATP-supported NAD+ replenishment helps explain the otherwise paradoxical restoration of cytosolic NAD+ and NAD+-dependent metabolism by a mitochondrial NAD+ importer. The requirement for glycolytic flux in SLC25A51-mediated protection further suggests that restored cytosolic metabolism is not simply a marker of healthier neurons, but instead is an active component of the protected state. Additional mechanisms, including mitochondrial SIRT3 activity or redox exchange through pathways such as the malate-aspartate shuttle, may also contribute to cytosolic recovery, but the present data identify ATP-supported cytosolic NAD+ resynthesis as a central mechanism by which mitochondrial NAD+ preservation sustains neuronal viability after SARM1 activation.

These findings place SARM1-mediated degeneration within an emerging view that mitochondrial NAD+ transport is an active determinant of cellular resilience rather than a passive metabolic reserve. Subcellular NAD+ pools are functionally interconnected, and mitochondrial NAD+ can buffer extra-mitochondrial NAD+ stress^25^. In other contexts, increasing SLC25A51-dependent mitochondrial NAD+ enhances liver regeneration and protects against cerebral ischemia/reperfusion injury^56,57^. Our study extends these observations to a distinct and mechanistically stringent context: persistent activation of an endogenous NADase. Under these conditions, SLC25A51 does not simply preserve mitochondrial function; it restores cytosolic NAD+-dependent metabolism and promotes stress-induced NAD+ synthesis despite ongoing SARM1 activity. Thus, SARM1 activation provides a stress test for compartmental NAD+coupling, revealing that maintenance of the mitochondrial NAD+ pool can support metabolic recovery outside the matrix even as cytosolic NAD+ is being actively consumed.

Our findings also intersect with recent work defining how SLC25A51 sustains mitochondrial NAD+ accumulation despite the electrochemical challenge of importing an anionic metabolite into the negatively charged matrix. SLC25 family carriers are often considered reversible and gradient-sensitive, yet mitochondrial NAD+ is maintained at a significantly higher concentration than cytosolic NAD+^58^. Cambronne and colleagues recently showed that mitochondrial membrane potential works together with charged residues in the SLC25A51 inner pore to enable sustained NAD+ import against its electrochemical gradient; dissipation of membrane potential or mutation of key pore residues promoted equilibration of matrix NAD+^59^. Our data extend this transport framework to a pathological stress state in which extra-mitochondrial NAD+ is actively consumed by SARM1. SLC25A51 overexpression increased mitochondrial NAD+ under basal conditions and preserved near-basal mitochondrial NAD+ during SARM1 activation. Importantly, SLC25A51 also preserved mitochondrial membrane potential, which is expected to support continued NAD+ import into the matrix^59^. Thus, mitochondrial NAD+ preservation and membrane potential maintenance may support a self-reinforcing protective state during SARM1 stress: preserved matrix NAD+ supports mitochondrial function, while preserved mitochondrial polarization supports continued SLC25A51-mediated NAD+ import. Defining how transporter abundance, carrier charge compensation, membrane potential, and ongoing NAD+ consumption shape net SLC25A51 flux during degeneration will be important, but is separable from our central conclusion: mitochondrial NAD+ availability governs metabolic resilience downstream of SARM1.

The *in vivo* data extend this model from acute pharmacologic SARM1 activation to chronic endogenous NAD+ stress. In the *Nmnat2*^V98M/R232Q^ model of a rare human genetic neuropathy^44^, SLC25A51 expression improved motor performance, axonal physiology, muscle preservation, and injury-associated Schwann cell responses, with more modest preservation of sciatic nerve axon density. The incomplete structural rescue in vivo suggests that mitochondrial NAD+ preservation raises the degenerative threshold rather than eliminating the upstream NMNAT2-driven, SARM1-dependent disease process. We employed a strong, constitutive promoter to overexpress SLC25A51, and therefore the protection cannot yet be attributed to a particular cell type. However, the degeneration in this model is driven by neuronal SARM1^44^, and together with the concordant protection of primary neurons *in vitro*, this supports neuronal mitochondrial NAD+ import as a likely *in vivo* mechanism of protection, although other cell types could also participate. The identification of a protective intervention downstream of SARM1 may be important therapeutically. Direct SARM1 inhibitors aim to block activation of the executioner enzyme, whereas enhancing mitochondrial NAD+ transport could preserve neuronal metabolism even when SARM1 is engaged, promoting resilience of metabolically stressed neurons. These approaches may therefore be complementary: SARM1 inhibition could reduce pathological NAD+ consumption, while mitochondrial NAD+ import could sustain ATP production, NAD+ resynthesis, and cytosolic metabolism in neurons that remain metabolically compromised.

## Acknowledgements

We thank the DiAntonio and Milbrandt lab members for technical support and insightful comments during this study, especially Xiaolu Sun, Kelli Simburger, Ashley Hollambey, Cassidy Menendez, and Sylvia Johnson. This work was supported by National Institutes of Health grant RF1AG097611 to A.D, J.M. and A.J.B., NS087632 to A.D. and J.M., and American Heart Association grant 24PRE1188417 to L.B. (2024). This work was also supported by the Hope Center Viral Vectors Core and the Needleman Center for Neurometabolism and Axonal Therapeutics at Washington University School of Medicine. This manuscript is the result of funding in whole or in part by the National Institutes of Health (NIH). It is subject to the NIH Public Access Policy. Through acceptance of this federal funding, NIH has been given a right to make this manuscript publicly available in PubMed Central upon the Official Date of Publication, as defined by NIH. The views expressed are those of the authors and do not necessarily represent the official views of the National Institutes of Health.

## Author contributions

L.B., Y.S., A.J.B., J.M. and A.D. conceived and designed the study. L.B., S.J.C., A.B., A.S., S.V.S., S.S. and H.K. performed the experiments. L.B. and Y.S. developed the methodology. L.B., S.J.C., A.B. and S.S. analyzed the data. L.B. and Y.S. curated the data. L.B. and A.D. wrote the original draft, and L.B., Y.S., A.J.B., J.M. and A.D. reviewed and edited the manuscript. Y.S., A.J.B., J.M. and A.D. supervised the study. L.B., A.J.B., J.M. and A.D. acquired funding.

## Competing interests

J.M. and A.D. are co-founders, scientific advisory board members, and shareholders of Disarm Therapeutics, a wholly owned subsidiary of Eli Lilly, and are scientific advisory board members of Asha Therapeutics and Cristae Therapeutics. L.B., J.M., and A.D. have a patent pending in relation to this work. The other authors declare no competing interests.

## Online Methods

### Animals

All animal experiments were performed in accordance with the policies and guidelines of the Institutional Animal Care and Use Committee (IACUC) of Washington University in St. Louis (protocol number 26-0043) and with the Guide for the Care and Use of Laboratory Animals of the National Institutes of Health. Mice were housed at no more than five per cage on a 12-h light/dark cycle with food and water available ad libitum. Timed-pregnant wild-type CD1 mice (Charles River Laboratories, Wilmington, MA) were used as the source of embryos for primary neuronal cultures unless otherwise indicated. *Nmnat3* knockout mice were previously derived from ES cells (Nmnat3^tm1(KOMP)Mbp^, Knockout Mouse Project (KOMP)) in our facility and crossed with C57BL/6 J mice for at least five generations^60^. These *Nmnat3* knockout and *Sarm1*-knockout mice were used for the corresponding primary embryonic cultures. Nmnat2^V98M/R232Q^ trans-heterozygous mice were maintained as described previously^44^. All embryos, with unknown sex, were used for *in vitro* experiments; for *in vivo* experiments, male and female mice were included, and different symbols were used to represent animals from each sex to visualize the distribution.

### Primary embryonic DRG cultures

Embryonic dorsal root ganglion (DRG) spot cultures were prepared essentially as previously described^12,39^. Tissue-culture plates were coated overnight at 37 °C with poly-D-lysine (10 mg/mL stock diluted 1:100 in water), washed three times with water, and then coated with laminin (Thermo-Fisher, Cat. #23017015; diluted 1:100 in water) for at least 4 h at 37 °C. DRGs were dissected from embryonic day 13.5–14.5 embryos, pelleted briefly at 800g for 4min, and incubated in 1 mL pre-warmed 0.05% trypsin for 13 min at 37 °C with gently flicking every ∼4-5 min. Trypsin was removed and ganglia were gently resuspended in 1 mL culture medium and triturated 20–30 times until no pellet remained. Cells were centrifuged at 800 × g for 4 min and resuspended in 60-100 µL of medium per embryo (∼3-7×10^6^ cells/ml). Cells were spotted at 2.5 µL per well of a 24-well plate or 1.2 µL per well of a 96-well plate, allowed to adhere for 13 min at 37 °C, and then gently topped with 500 µL (24-well) or 100 µL (96-well) of medium per well.

Culture medium consisted of Neurobasal medium (Thermo-Fisher, Cat. #21103049) supplemented with L-glutamine, penicillin–streptomycin, 2% B27 (Thermo-Fisher, Cat. #17504044), 50 ng/mL nerve growth factor (Envigo Bioproducts, Cat. #B5017), and 1 µM 5-fluoro-2′-deoxyuridine (FDU; Sigma-Aldrich, Cat. #F0503). On DIV4, half of the medium was removed and replaced with fresh medium. All experiments were performed at DIV6–8 unless otherwise stated.

### Primary embryonic cortical neuron cultures

Tissue-culture plates were coated overnight at 37 °C with poly-D-lysine (10 mg/mL stock diluted 1:100 in water), then washed three times with water before plating. Cortices were dissected from embryonic day 13.5-14.5 embryos and meninges were peeled off. Cortices were resuspended in final culture media volume for plating (12mL for every 2-3 embryos), and tissue was dissociated without enzymatic digestion by triturating 10 times using a 10mL pipette, then filtered using a 70μm cell strainer. The cell suspension was plated at 500 µL (24-well) or 100 µL (96-well) per well, from a ∼200-400k/mL suspension. Culture medium consisted of Neurobasal medium (Thermo-Fisher, Cat. #21103049) supplemented with L-glutamine, penicillin–streptomycin, 2% B27 (Thermo-Fisher, Cat. #17504044), and 1 µM 5-fluoro-2′-deoxyuridine (FDU; Sigma-Aldrich, Cat. #F0503). On DIV4, half of the medium was removed and replaced with fresh medium. All experiments were performed at DIV6–8 unless otherwise stated.

### Plasmids

Lentiviral transfer plasmids were based on the FUGW backbone (Addgene #14883)^61^. Constructs encoded N-terminal 3x-FLAG-tagged versions of human SLC25A51, the transport-dead point mutant SLC25A51-K91A, and the yeast mitochondrial NAD+ carrier Ndt1, and GFP or empty vector were used as controls. For *in vivo* work, the 3x-FLAG-SLC25A51 construct was cloned into a different backbone for AAV expression; the AAV genome comprised the human EF-1α promoter driving 3×FLAG-tagged SLC25A51, followed by WPRE and a human growth hormone polyadenylation signal, flanked by AAV2 inverted terminal repeats. The ratiometric NAD+ sensor FiNad^34^ (gift from Philip Williams) was targeted to the cytosol or to the mitochondrial matrix using the COX8A mitochondrial targeting sequence and cloned into the same FUGW backbone. The ATP sensor iATPSnFR2 targeted to the cytosol or to the mitochondrial matrix (Addgene 236407 and 209723), was also cloned into the FUGW backbone under the EF-1a promoter, which replaced the UbC promoter. All plasmids were verified by whole-plasmid long-read sequencing (Plasmidsaurus). Plasmids generated in this study are available from the corresponding author on request.

### Lentiviral transduction

Lentivirus was produced by transfecting HEK293T cells maintained in DMEM supplemented with L-glutamine, 10% FBS, and penicillin–streptomycin. One day before transfection, cells were passaged with 0.05% trypsin, collected by centrifugation at 400 × g for 3 min, and seeded into 6-well plates; medium was replaced with 2 mL fresh medium per well immediately before transfection. For each well, 100 µL Opti-MEM and 6 µL FuGENE 6 Transfection Reagent (Promega, Cat. #E2691) were combined and incubated for at least 5 min at room temperature. In parallel, 200 ng pCMV-VSV-G (Addgene, Cat. #8454), 600 ng psPAX2 (Addgene, Cat. #12260), and 200 ng of the transfer plasmid were combined per well. The Opti-MEM/FuGENE mixture was added to the DNA, incubated for 15 min, and added dropwise to the cells. Forty-eight hours after transfection, virus-containing medium was collected and centrifuged at 4,000 × g for 5 min at 4 °C to pellet residual cells, and the supernatant was transferred to fresh tubes. Virus was concentrated approximately 10-fold by adding Lenti-X Concentrator (Takara Bio, Cat. #631232) at one-third of the medium volume, mixing by inversion, incubating on ice for 30 min to overnight, and centrifuging at 1,500 × g for 45 min at 4 °C; pellets were resuspended in one-tenth of the original volume with Neurobasal. Concentrated virus was aliquoted and stored at −80 °C. All materials contacting lentivirus were decontaminated with bleach before disposal. Neurons were transduced by adding virus directly to the culture medium at DIV0–2 at a final concentration of 2.5-5% in volume. Neurons transduced for SLC25A51 were compared to control neurons transduced with either empty vector or GFP.

### Lentiviral shRNA knockdown

Slc25a51 was knocked down with MISSION lentiviral shRNA constructs (Sigma-Aldrich; TRCN0000123900) supplied as bacterial glycerol stocks, together with a non-targeting shRNA control (Sigma-Aldrich; SHC002). Plasmid DNA was prepared from the glycerol stocks and lentivirus was produced and applied as described above. One day after plating, cortical and DRG neurons were co-transduced with lentivirus expressing human BCL-XL (NP_612815) at 1% of the culture volume and shRNA lentivirus at 1.5% of the culture volume, and were then cultured for a further week before analysis. Knockdown efficiency was determined by quantitative RT–PCR on sister cultures as described below. For rescue experiments, an SLC25A51 construct was confirmed to be shRNA-resistant and delivered by lentivirus alongside the shRNA (2.5% in volume).

### Quantitative RT–PCR

Total RNA was isolated from cortical neurons one week after transduction using the SV Total RNA Isolation System (Promega; Cat. #Z3505). Complementary DNA was synthesized with the iScript cDNA Synthesis Kit (Bio-Rad; Cat. #1708891) in 20 µL reactions containing 4 µL 5× iScript reaction mix, 1 µL iScript reverse transcriptase, template RNA and nuclease-free water. RNA input was normalized across all samples within a batch to the lowest concentration available in that batch. Quantitative PCR was performed with TaqMan Fast Advanced Master Mix (Thermo Fisher Scientific; Cat. #4444557) and TaqMan gene expression assays for Slc25a51 (assay ID Mm05911874_g1) and Actb (assay ID Mm02619580_g1) in 20 µL reactions. Each sample was assayed in three technical replicates, which were averaged before analysis. Slc25a51 expression was normalized to Actb and expressed relative to the mean of the non-targeting shRNA samples by the 2^−ΔΔCt^ method.

### Drug treatments

SARM1 was activated with vacor (Chem Service N-13738), dissolved in DMSO at 50 mM for storage at −80 or −20°C and used at 2.5 or 5 µM for cortical neurons and 10 or 12.5 µM for DRG neurons, as indicated in the figure legends. An equal volume of DMSO served as vehicle control in all experiments. The following compounds were also used: the SARM1 inhibitor NB-7^20^ (3 µM); the GAPDH inhibitor heptelidic acid (500 nM; Cayman Chemical Company 14079); the ATP synthase inhibitor oligomycin (250 nM; Sigma-Aldrich 75351); the NAMPT inhibitor FK866 (10 nM; Sigma-Aldrich F8557); nicotinamide riboside (NR; 1 mM; gift from ChromaDex); and deuterium-labeled nicotinamide (300 µM; nicotinamide-2,4,5,6-d4, D4-Nam; C/D/N Isotopes, Pointe-Claire, QC).

### Analysis of axon degeneration

Axon degeneration was quantified using the degeneration index (DI) as described previously^3,12,39^. DRG spot cultures in 96-well plates were treated with vacor or, for axotomy experiments, transected at DIV6-8 with a microsurgical blade under a dissecting microscope. Bright-field images of distal axons (10 fields per well) were acquired at the indicated times after treatment or axotomy on a high-content imager (Operetta, PerkinElmer, Waltham, MA) with a 20× objective. The DI was calculated in ImageJ (NIH) from images of the same fields at each time point, using our previously published ImageJ macro with higher values indicating greater fragmentation^62^. At least three independent experiments were performed for each condition.

### Cell death

Neuronal viability was assessed 24 h after vacor treatment. DRG cultures were stained with 1:1,000 propidium iodide (PI) and 1:10,000 Hoechst 33342 (1ug/mL; InvitrogenH3570), and images were acquired on a Molecular Devices IN Cell Analyzer at 20x, with 9 images per well. DRG cell death was quantified as the fraction of nuclei that were PI-positive within the Hoechst-positive population using the IN Carta software (Molecular Devices). Cortical neuron cultures were incubated with 1:2,000 NeuroFluor™ NeuO (Stem Cell Technologies 01801) and imaged on a Molecular Devices IN Cell Analyzer at 20x, with 5-6 images per well. Total green intensity was used to quantify viable cells using the IN Carta software (Molecular Devices).

### Flow cytometry

Cortical neurons in 96-well plates were dissociated by removing the medium and adding 50 µL per well of pre-warmed 0.05% trypsin, followed by 5 min at 37 °C. Trypsin was diluted with 200 µL of medium per well, cells were triturated 8–10 times, and the suspension was transferred to a round-bottom plate. Cells were centrifuged at 400 × g for 5-10 min, the supernatant was decanted, and pellets were resuspended in flow buffer (filtered DPBS with 1% BSA) containing Hoechst 33342 at 1:10,000. Suspensions were triturated 8–10 times and filtered through a 70 µm filter into a second round-bottom plate immediately before acquisition. Unstained cells and single-color controls were included, and Hoechst plus propidium iodide staining was used as a positive control for gating out dead cells. Data were acquired on CytoFLEX S Flow Cytometer (Beckman Coulter) and analyzed in FlowJo (BD Biosciences); the BP450 (Hoechst), FITC (green), and ECD (mCherry) channels were recorded. Debris was excluded using SSC-A/FSC-A, singlets were selected using FSC-W/FSC-H, and viable cells were identified as the Hoechst-low population (example sample showing gating strategy in Supplementary Information). The Hoechst-high population overlapped with PI positivity in untransduced cells, supporting its identification as dead or membrane-compromised cells. Sensor-positive cells were subsequently gated using red-channel fluorescence. Flow cytometry was performed on cortical neuron cultures only, as DRG neurons were grown as spot cultures, which yield a low cell count and too much debris for flow cytometric analysis.

### NAD+ sensor measurements

Compartment-specific NAD+ levels were monitored in live neurons using the ratiometric fluorescent NAD+ sensor FiNad^34^ targeted to the cytosol or to the mitochondrial matrix. Neurons were transduced with sensor-encoding lentivirus at DIV0–2 and analyzed at DIV5-7. Sensor localization was validated by confocal imaging (Extended Data Fig. 1c). Sensor responsiveness was validated pharmacologically by treatment with the NAMPT inhibitor FK866 and the NAD+ precursor nicotinamide riboside. For quantification, cells were prepared for flow cytometry as described above and the green:red emission ratio was recorded for each cell that was positive for the sensor/mCherry; the population ratio was normalized to the DMSO-treated, empty-vector control within each experiment. Because the FiNad sensor operates over a limited dynamic range, modest changes in normalized ratio correspond to substantially larger changes in absolute NAD+ concentration^34^.

### ATP sensor measurements

Cytosolic and mitochondrial ATP were monitored in live neurons expressing the corresponding iATPSnFR2 sensor^36^. Neurons were transduced with sensor-encoding lentivirus at DIV0–2 and analyzed at DIV6-8. For timecourses, neurons were imaged on the ImageXpress HCS at 40x under environmental control (37 °C, 5% CO2) with images acquired every 20 min for 10 hours following drug treatment, with multiple fields per well. Neurons were otherwise imaged on the IN Cell Analyzer (Molecular Devices) at 20x. Total green fluorescence intensity per field was measured on the IN Carta Image Analysis Software and normalized to the t = 0 h value and to the DMSO-treated control.

### Mitochondrial membrane potential (TMRM)

Mitochondrial membrane potential was assessed with tetramethylrhodamine methyl ester (TMRM; 100 nM; ThermoFisher T668). Cortical neurons transduced with SLC25A51 or control virus were treated with vacor (5 µM) or DMSO for 4 h, loaded with TMRM, and imaged on the IN Cell Analyzer (Molecular Devices) at 20x. Total TMRM intensity was quantified on the IN Carta Image Analysis Software and expressed relative to the DMSO-treated control.

### Reactive oxygen species

The presence of reactive oxygen species was assessed in primary mouse DRG neuron cultures using CM-H DCFDA (Invitrogen C6827). Cultures were incubated with 10 μM CM-H DCFDA for 30 min before baseline imaging, and the dye remained in the culture medium throughout the experiment. Fluorescence images were acquired at 0 and 6 h using an IN Cell Analyzer 6500 HS. Mean cellular CM-H DCFDA fluorescence intensity was quantified using IN Carta image-analysis software, and the 6 h intensity was normalized to the corresponding 0 h value.

### Subcellular fractionation

Cytosolic and mitochondrial fractions were prepared from cultured neurons by differential centrifugation, adapted from the Abcam mitochondrial purification protocol for western blot samples. Buffers were prepared as follows: NKM buffer (1 mM Tris-HCl, pH 7.4, 0.13 M NaCl, 5 mM KCl, 7.5 mM MgCl2); homogenization buffer (10 mM Tris-HCl, 10 mM KCl, 0.15 mM MgCl2, 1 mM PMSF, 1 mM DTT); and 2 M sucrose. All buffers were ice-cold and all steps were performed at 4 °C or on ice. Cells were trypsinized with 0.05% trypsin for 5 min at 37 °C in half the feeding volume, diluted at least 1:5 in culture medium, and collected by centrifugation at approximately 370 × g for 10 min. The supernatant was decanted and cells were resuspended in 10 packed cell volumes of NKM buffer (1 packed cell volume = 100 µL); cells were re-pelleted and this wash was performed three times in total. Cells were then resuspended in 6 packed cell volumes of homogenization buffer, transferred to a glass homogenizer, and incubated for 10 min on ice before homogenization with a tight-fitting pestle, for approximately 30 strokes. The homogenate was poured into a tube containing 1 packed cell volume of 2 M sucrose and mixed gently by inversion. Unbroken cells, nuclei, and large debris were pelleted at 1,200 × g for 5 min and the supernatant was transferred to a fresh tube; this clearing step was repeated twice in total, discarding the pellet each time and transferring an identical volume across all samples. Mitochondria were then pelleted at 7,000 × g for 10 min. The resulting supernatant, the cytosolic fraction, was re-centrifuged at 7,000 × g for 10 min to remove residual mitochondria and the cleared supernatant was transferred to a fresh tube for immunoblotting or metabolite extraction. The mitochondrial pellet was washed in 3 packed cell volumes of cold DPBS, or ice-cold 0.9% NaCl when the fraction was destined for metabolite analysis, re-pelleted at 9,500 × g for 5 min, and either resuspended in 100 µL of protein lysis buffer or processed for metabolite extraction as described below. Fraction purity was confirmed by immunoblotting for the cytosolic marker HSP90 and the mitochondrial marker COX4.

### Metabolite collection and measurement

Metabolites were extracted from adherent cultures approximately one week after plating. Ice-cold 0.9% NaCl (phosphate-free) and ice-cold 50% LC/MS-grade methanol in Milli-Q water were prepared in advance and chilled on ice. Plates were placed on ice, the medium was removed as completely as possible, and ice-cold 0.9% NaCl was added gently without disturbing the cells (1 mL per well of a 24-well plate well). The NaCl solution was then removed and replaced with 50% methanol/water (160 µL per well of a 24-well plate), and plates were incubated on ice for 5 min. The extract (160 µL) was collected into PCR tube strips without disturbing the cell layer. One-third volume of chloroform (approximately 50 µL) was added, samples were shaken vigorously and centrifuged at 20,380 × g for 10 min at 4 °C, and the aqueous phase (typically 140 µL per 24-well sample) was collected, lyophilized, and stored at −80 °C until analysis. For protein normalization, 100 µL of 0.1% SDS was added to the remaining cell material in each well and incubated at room temperature for 15–30 min. The lysate was collected and protein concentration was determined with a Pierce BCA Protein Assay kit (Thermo-Fisher, Cat. #23227) with standards run in triplicates and samples run in duplicates; protein concentration per sample was determined from the mean of the replicates. DRG neuron metabolite measurements were normalized well-by-well to protein concentration. Cortical neuron metabolite measurements were not normalized, as cell counts were constant across wells.

For tissue metabolites, sciatic nerves were dissected, immediately flash-frozen after peeling the epineurium, and stored at −80 °C. Extraction was performed on frozen tissue without thawing, with samples kept on dry ice throughout handling. Because sciatic nerves are too small to weigh accurately, 160 µL of ice-cold 50% LC/MS-grade methanol in water pre-chilled on ice was added into the 1.5 mL microcentrifuge tubes containing the collected nerves; these samples were not weighed and were instead normalized to protein as described below. Samples were allowed to settle to the bottom of the tube and were homogenized by sonication (output 1.5, 50% duty cycle, 10-20 strokes) with the probe tip positioned just above the tissue without contacting it.

Homogenization was judged complete when the homogenate was uniformly cloudy across all samples; residual nerve sheath material remained and was not further disrupted. Homogenates were centrifuged at 20,380 × g for 10 min at 4 °C and the entire supernatant was transferred to a fresh tube. One-third volume of chloroform (approximately 50 µL) was added, samples were shaken vigorously and centrifuged at maximum speed for 10 min, and the aqueous phase was collected. A second chloroform extraction was performed with a further 50 µL of chloroform, followed by vigorous shaking and centrifugation at maximum speed for 15 min; 120 µL of the aqueous phase was transferred to a fresh tube, and when less than this volume was recovered the collected volume was recorded and carried through the calculation. Aqueous extracts were lyophilized and stored at −80 °C until analysis. For protein normalization, the pellet remaining after the initial clearing spin was retained, kept on dry ice, and stored at −80 °C until assay; 500 µL of 1% SDS in water was added, samples were sonicated (output 2.5, 50% duty cycle, 10 strokes) and centrifuged at 15,000 × g for 5 min, protein concentration was determined using a Pierce BCA Protein Assay kit. Tissue metabolite levels were normalized to protein content on a per-sample basis.

On the day of measurement, lyophilized samples were reconstituted in 5 mM ammonium formate and centrifuged at 13,000 × g for 10 min at 4 °C, before 10 µL of clear supernatant was injected. NAD+, NMN, and cADPR were measured by LC-MS/MS as described previously^12,41,63^. Samples were separated on a C18 reverse-phase column (Atlantis T3, 2.1 × 150 mm, 3 µm; Waters, Milford, MA) using an Agilent 1290 Infinity LC at a flow rate of 0.15 mL/min, with 5 mM ammonium formate as mobile phase A and 100% methanol as mobile phase B. Metabolites were eluted with the following gradient: 0–10 min, 0–70% B; 10–15 min, 70% B; 16–20 min, 0% B. Detection used an Agilent 6470 triple-quadrupole mass spectrometer (MassHunter) in positive-ion electrospray multiple reaction monitoring (MRM) mode with compound-specific parameters: NAD+ (Sigma, Cat. #N1636) 664 > 428, fragmentor (F) = 160 V, collision energy (C) = 22 V, cell accelerator (CA) = 7 V; NMN (Sigma, Cat. #N3501) 335 > 123, F = 135 V, C = 8 V, CA = 7 V; cADPR (Sigma, Cat. #C7344) 542 > 428, F = 100 V, C = 20 V, CA = 3 V. Serial dilutions of authentic standards in 5 mM ammonium formate were used for calibration. Metabolites were quantified against standard curves using the MassHunter quantitative analysis tool (Agilent) and normalized to protein concentration. Broad targeted metabolomic profiling covering 218 metabolites spanning glycolysis, the pentose phosphate pathway, the TCA cycle, and purine metabolism was performed as instructed (Metabolomics dMRM Database and Method, Agilent Technologies^64^).

Metabolites were extracted as described above and chromatographically separated using a reverse phase column (Zorbax RRHD Extend-C18, 2.1 × 150 mm, 1.8 μm, Agilent Technologies) with mobile buffer A (3% MeOH, 10 mM tributylamine, 15 mM acetic acid) and 0%–99% gradient of buffer B (100% MeOH, 10 mM tributylamine, 15 mM acetic acid). Metabolites were analyzed by mass spectrometer (6470, Agilent Technologies) using dynamic multiple reaction monitoring. The compound signatures, including retention time and qualifying ions, are predetermined in the dynamic multiple reaction monitoring database and data analysis tool automatically detects these compounds. After filtering out low abundance metabolites (area under curve <200), metabolites with missing values in more than 50% of samples were excluded, and remaining missing values were replaced with one fifth of the minimum positive value for that metabolite. The relative amount of each metabolite was calculated and normalized to protein as specified above. For the targeted metabolomic dataset, principal component analysis was performed on 63 reliably measured metabolites following blank subtraction, protein normalization, and log transformation on MetaboAnalyst 6.0^65^; group separation was tested by pairwise PERMANOVA with 999 permutations, and p values were adjusted for multiple comparisons by the Benjamini–Hochberg method; ellipses on PCA plots denote 95% confidence intervals.

### D4-NAD+ synthesis assay

NAD+ synthesis was measured by stable-isotope labelling as described previously^12,39^. Neurons were topped off with D4-nicotinamide (nicotinamide-2,4,5,6-d4) at a final concentration of 300 µM ∼2.5 hours after vacor vs DMSO treatment, and metabolites were collected 2.5 h later as described above. Labeled (D4-NAD+; heavy) NAD+ was quantified by LC-MS/MS. Newly synthesized NAD+ was measured as D4-NAD+, which had to have been synthesized within the previous 2.5 hours.

### Glycolytic flux assay

Neurons were treated with vacor for 5 hours, then media was removed and cells were washed once with ice-cold PBS. The media was then replaced with a glucose-free and pyruvate-free version with 1 mM D-Glucose-^13^C_6_ (Sigma-Aldrich 389374) and placed back in incubator for 3 hours before metabolite collection.

### Protein lysis

For cultured cells, lysis buffer was prepared fresh or within a day of protein collection, as follows: 102 µL RIPA buffer (Sigma-Aldrich R0278), 17 µL 7× protease inhibitor (from tablet, one tablet in 1.5 mL water), and 1.2 µL of each of two phosphatase inhibitors per ∼120 µL of final buffer. Medium was aspirated, wells were washed with cold DPBS, and lysis buffer was added at 100 µL per well of a 24-well plate or per mitochondrial pellet after subcellular fractionation. Cells were scraped with a pipette tip or by triturating a little more harshly, collected, and the same volume was carried to a second well when pooling. Lysates were centrifuged at 2,000 x g for 4 min at 4 °C, and the supernatant was transferred to a fresh tube and stored at −80 °C. For tissue, samples were homogenized by sonication in ∼100 µL of lysis buffer (output 2.5, 50% duty cycle, 10-20 strokes). Lysates were centrifuged at 17,000 × g for 15 min at 4 °C and the supernatant was collected for BCA assay and immunoblotting. Sciatic nerves were flash-frozen immediately after dissection and kept on dry ice until lysis.

### Western blot

Protein concentration was determined with a Pierce BCA Protein Assay kit against a BSA standard curve (25–2,000 µg/mL) read at 562 nm on a plate reader, and sample concentrations were interpolated from the linear fit. Equal protein amounts per lane, depending on lysate type and target, were combined with 6× Laemmli buffer and 1× RIPA buffer (Sigma-Aldrich R0278) to a constant volume, heated at 95 °C for 10 min, and returned to ice. Samples were resolved on 4–20% Mini-PROTEAN TGX precast gels (Bio-Rad) in Tris-glycine running buffer (25 mM Tris, 192 mM glycine, 0.1% SDS) at 40 V for 10-20 min, until samples fully migrated into the gel, followed by 100 V until the dye front reached the bottom of the gel, with Precision Plus Dual Color ladder in the flanking lanes. Proteins were transferred to nitrocellulose using Trans-Blot Turbo Transfer Packs (Bio-Rad) on the TURBO setting (7 min). Membranes were blocked for 0.5-2 hours at room temperature in 5% non-fat dry milk in TBS with 0.2% Tween-20 (TBST), incubated with primary antibody overnight at 4 °C, washed three times for 5-10 mins each in TBST, incubated with HRP-conjugated secondary antibody (1:5,000) for 1-2 hours at room temperature, and washed three more times for 5-10 mins each in TBST. Signal was developed with ECL substrate (approximately 2mL per membrane) and imaged on Syngene G:BOX with exposure times between 3s and 30s. Band intensities were quantified in ImageJ using the gel-analysis tool; the protein of interest was normalized to its loading control within each lane, and these ratios were then normalized to the mean of the control samples on the same blot. The following antibodies were used: anti-HSP90 (1:2000), anti-COX4 (1:1000), anti-c-Jun (1:1000), anti-cofilin (1:1000), and anti-FLAG(1:1000) together with Peroxidase AffiniPure Goat Anti-Rabbit IgG and Goat Anti-Mouse IgG (1:5,000; Jackson ImmunoResearch, Cat. #111-035-045 and #115-035-003).

### AAV production and delivery

3x-FLAG-SLC25A51 or GFP was expressed from the EF1α promoter in an AAV genome and packaged as a dual AAV9/PHP.eB/S capsid preparation produced by the Hope Center Viral Vectors Core at Washington University School of Medicine. 20 µL AAV aliquots (3.7-4×10^13^vg/mL) were diluted with 80 µL sterile saline (per animal).

Nmnat2^V98M/R232Q^ trans-heterozygous mice were randomly assigned to experimental groups and injected intrathecally, between postnatal days 13 and 19, with the full amount of 100 µL at around L5-6, while under light anesthesia (avertin). Prior to injection, mice were re-genotyped for confirmation and transgene expression was confirmed in spinal cord and sciatic nerve at endpoint by FLAG immunoblotting. Male and female mice were both tested and plotted using different symbols (M: squares; F: circles). Investigators were blinded to treatment group for all behavioral and histological analyses.

### Inverted screen

Muscle strength was assessed by the inverted screen test at 2, 3, 4, 5, and 6 months of age. Each mouse was placed on a wire mesh screen, which was then inverted, and the latency to fall was recorded with a maximum trial duration of 2 min. Each mouse underwent three trials separated by ≥5-min rest periods, and the mean of the three trials was used as that animal’s value. If any two trials had more than 20 seconds of difference, a fourth trial was run and included in the mean for that mouse. If a mouse fell while trying to reach over the edge of the inverted screen, the mouse was rapidly placed back on the inverted screen and given one more chance to finish the trial while subtracting 10 seconds from the latency to fall.

### Nerve electrophysiology

Compound muscle action potentials (CMAPs) were recorded using a Nicolet VikingQuest electromyography system as described previously^8^. Mice were anaesthetized with isoflurane in oxygen delivered by nose cone, anesthetic depth was monitored by toe pinch throughout, and ophthalmic ointment was applied. Animals were maintained on a heating pad between recordings; the pad was switched off during acquisition to avoid electrical interference. A subdermal recording electrode was placed in the plantar surface of the foot and a stimulating electrode was placed subcutaneously at the ankle and then at the sciatic notch. Stimulation was performed at 2.45mA as supramaximal current. Amplitude was recorded from both the left and right sides of each animal and averaged.

### Muscle atrophy

At 6 months of age, tibialis anterior (TA) muscles were dissected bilaterally, trimmed of tendon and connective tissue, and weighed on an analytical balance. The two values were averaged and the average TA muscle mass per mouse was normalized to whole-body weight measured immediately before euthanasia.

### Nerve structural analysis

Sciatic nerves were processed for semithin sectioning as described previously^8,66^. All solutions were prepared in Millipore-distilled water. Nerves were carefully dissected without stretching or clamping except at the ends, laid on a wooden stick to keep them straight, and immersion-fixed in 3% glutaraldehyde (Polysciences, cat. no. 01909) in 0.1 M Sorensen’s phosphate buffer (PB), pH 7.4, for at least 24 h at 4 °C. Nerves were then removed from the stick, washed three times in 0.1 M PB, and post-fixed in 1% osmium tetroxide in 0.1 M PB (250 µL per nerve) overnight at 4 °C on a rocker. After osmication, nerves were washed three times in 0.1 M PB. Samples were dehydrated through a graded ethanol series (3 × 50%, 3 × 70%, 3 × 95%, 3 × 100%; 10–15 min each), transferred through a 1:1 mixture of propylene oxide and 100% ethanol, and then through 100% propylene oxide twice (10–15 min each; longer incubations were avoided to prevent tissue damage). Nerves were infiltrated with Araldite resin/propylene oxide at 50:50 overnight at 4 °C, then at 70:30 and 90:10, each for several hours at room temperature followed by overnight at 4 °C, and finally in 100% Araldite for approximately 2 h at room temperature. Araldite resin was prepared by mixing Araldite 502 (Polysciences, cat. no. 00552), dodecenylsuccinic anhydride (DDSA; Polysciences, cat. no. 00563), and DMP-30 (Polysciences, cat. no. 00553) at 12:9:1. Nerves were embedded in plastic molds with paper labels and the resin was polymerized overnight at 60 °C. Semithin sections of 400–600 nm were cut on a Leica EM UC7 ultramicrotome, mounted on slides, and stained with toluidine blue. Images were acquired on a Leica Thunder Imager and axon density was quantified in MyelTracer^67^: two 500×500μm² cropped sections from motor-predominant areas (larger axons) were used for axon counts and averaged per mouse; axons along the image edges were counted if ∼>70% was within the cropped image. All quantification was performed blinded to genotype and treatment.

### Statistical analysis

Statistical analyses were performed in GraphPad Prism 11. For *in vitro* experiments, the experimental unit was an independently treated culture well. Neurons from multiple embryos of a litter were pooled before plating to generate a common neuronal suspension, which was distributed among wells and independently assigned to experimental conditions. For *in vitro* imaging experiments, multiple image fields within each well were treated as technical subsamples and averaged to generate one value per well. Experiments were repeated using independently prepared neuronal cultures from separate dissections on at least three experimental days to ensure reproducibility across culture preparations. Statistical tests used are stated in each figure legend.

Comparisons involving two factors were analysed by two-way ANOVA with an interaction term; time-course data were analysed by two-way repeated-measures ANOVA matched by row. Mixed-effects models were used for repeated-measures time courses with missing values. Pairwise comparisons between two conditions used two-tailed unpaired t tests or Welch’s t-test. Multiple comparisons following ANOVA were corrected by Tukey’s test for all-versus-all comparisons, Šídák or Holm-Šídák for preselected comparisons. Significance thresholds are indicated as *p < 0.05, **p < 0.01, ***p < 0.001, ****p < 0.0001. For the targeted metabolomic dataset, principal component analysis was performed on 63 reliably measured metabolites following blank subtraction, normalization to protein concentration, and log transformation on MetaboAnalyst 6.0; group separation was tested by pairwise PERMANOVA with 999 permutations, and p values were adjusted for multiple comparisons by the Benjamini– Hochberg method; ellipses on PCA plots denote 95% confidence intervals.

### Reporting Summary

Further information on research design is available in the Nature Portfolio Reporting Summary linked to this article.

### Data Availability

Upon publication, all data will be freely available from the Digital Commons Data @ Becker Repository [doi, url]. Source data will be provided with this paper.

## Extended Data figures

**Extended Data Fig. 1:**
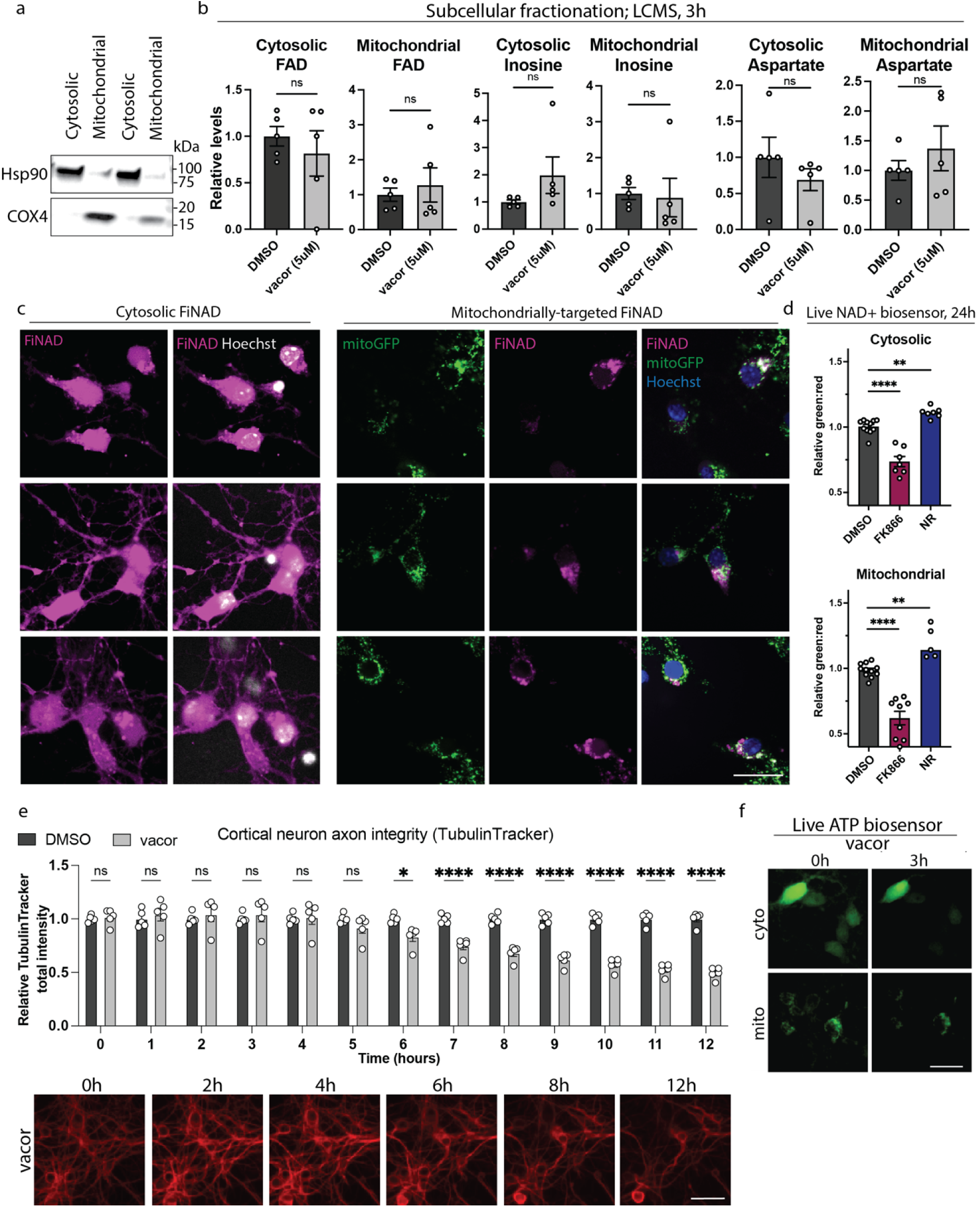
a. Western blot confirmation that the cytosolic and mitochondrial fraction following subcellular fractionation were enriched with cytosolic protein Hsp90 and mitochondrial protein Cox4, respectively. b. Cytosolic and mitochondrial FAD, inosine, and aspartate levels in wild-type primary mouse cortical neurons treated with vacor (5 µM) vs DMSO for 3 hours. Metabolites are shown relative to DMSO-treated control, as measured by LC-MS following subcellular fractionation. Error bars show mean ± SEM, with data points representing independently treated culture wells (n=5) pooled across 3 independent culture preparations. Statistical significance was determined with a two-tailed unpaired t-test. c. Representative images showing FiNad sensor (magenta) localization: cytosolic sensor was diffuse while the mitochondrial sensor co-localized with mitoGFP (green). Hoechst is shown in grey/blue. Scale bar, 20 µm. d. Cytosolic and mitochondrial measurements of live NAD+ sensor FiNad in primary cortical neurons at 24 hours following FK866 (10 nM) or nicotinamide riboside (1 mM) treatment. Green:red ratio was measured with flow cytometry, and is shown relative to the ratio in DMSO-treated control. Error bars show mean ± SEM, with data points representing independently treated culture wells (n=5-11) pooled across 3 independent culture preparations. Statistical significance was determined with an ordinary one-way ANOVA. e. Total TubulinTracker intensity relative to DMSO control, imaged every hour following treatment of cortical neurons with 2.5 µM vacor vs control, with representative images. Error bars show mean ± SEM, with data points representing independently treated culture wells (n=5). Statistical significance was determined by 2-way ANOVA with Holm-Šídák’s multiple comparisons test. Scale bar, 25 µm. f. Representative images of live cytosolic (cyto) and mitochondrial (mito) ATP sensor (green) at 0 and 3 hours following treatment with 2.5 µM vacor in cortical neurons. Scale bar, 25 µm.

**Extended Data Fig. 2:**
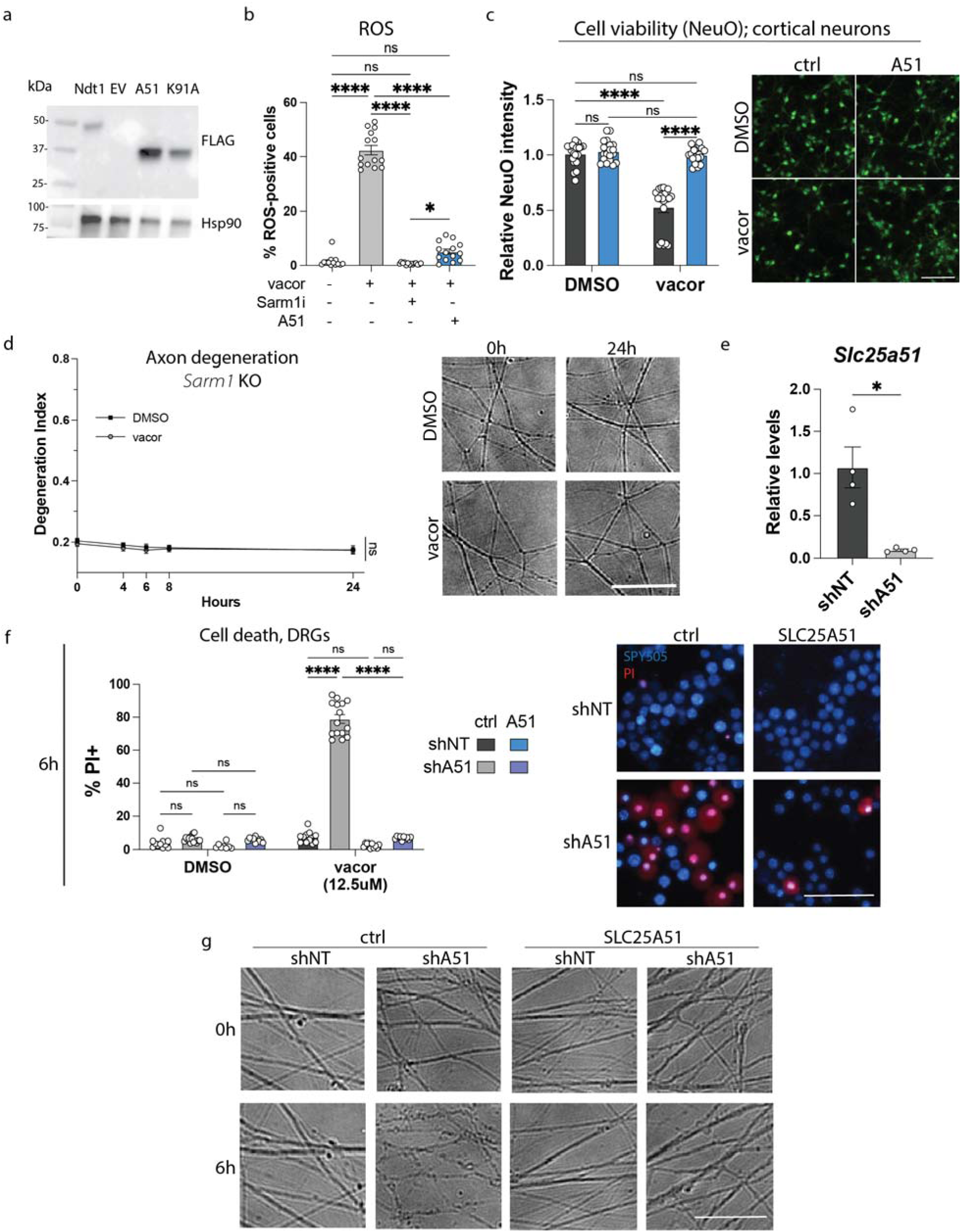
a. Western blot images of cortical neuron protein lysates one week after transduction with empty vector (EV) control, 3x-FLAG-SLC25A51 (A51), 3x-FLAG-Ndt1 (Ndt1), or 3x-FLAG-SLC25A51-K91A (K91A). Hsp90 was used as a loading control and FLAG bands were observed in the Ndt1, A51, and K91A conditions, validating appropriate construct expression. b. Percent of primary DRG neurons positive for reactive oxygen species based on quantification of microscopy images of the oxidative stress indicator CM-H2DCFDA after treatment with vacor (10 µM) ± SARM1 inhibitor NB-7(3 µM) or A51 lentiviral overexpression. Error bars show mean ± SEM, with data points representing independently treated culture wells (n=13-14) pooled across 3 independent culture preparations. Statistical significance was determined by one-way ANOVA with Tukey’s multiple comparisons test. c. Cell viability in primary cortical neurons transduced with A51 vs control (GFP/empty vector) and treated with vacor (2.5 µM) for 24 hours. Error bars show mean ± SEM, with data points representing independently treated culture wells (n=20) pooled across 3 independent culture preparations. Statistical significance was determined by 2-way ANOVA with Tukey’s multiple comparisons test. Representative images of live neuronal stain NeuO (green) are shown at 24 hours following treatment with 2.5 µM vacor in control (empty vector) vs SLC25A51-overexpressing cortical neurons. Scale bar, 100 µm. d. Axon degeneration quantification with representative images in primary *Sarm1*-knockout DRG neurons treated with vacor (12.5 µM) vs DMSO over 24 hours. Degeneration index was quantified, with an increase indicating increased degeneration. Error bars show mean of independently treated culture wells ± SEM, with n=5. Statistical significance was determined by a 2-way ANOVA with Tukey’s multiple comparisons test with pairwise comparisons shown for 24h. Scale bar, 50 µm. e. *Slc25a51*expression levels, normalized to *actb*, as measured by qRT-PCR one week after cortical neuron transduction with non-targeting shRNA vs *Slc25a51*-targeting shRNA. Error bars show mean of independently treated culture wells ± SEM, with n=4. Statistical significance was determined with Welch’s t-test. f. Cell death in primary DRG neurons transduced with non-targeting shRNA (shNT) or *Slc25a51-*targeting shRNA (shA51), in addition to shRNA-resistant SLC25A51 overexpression or control (empty vector), and treated with vacor (12.5 µM) for 6 hours. Cell death was measured by the fraction of nuclei (SPY505+) positive for propidium iodide (PI). Error bars show mean ± SEM, with data points representing independently treated culture wells (n=15) pooled across 3 independent culture preparations. Statistical significance was determined with a 2-way ANOVA with Holm-Šídák’s multiple comparisons test. Representative images showing SPY505+ nuclei (blue) and propidium iodide (PI) positive dead cells (red) at 6 hours post vacor. Scale bar, 50 µm. g. Representative axon images of primary DRG neurons transduced with shRNA-resistant A51 or control (empty vector), in addition to non-targeting shRNA (shNT) or *Slc25a51-*targeting shRNA (shA51), and treated with vacor (12.5 µM) vs DMSO; shown at 0 and 6 hours post-vacor. Scale bar, 50 µm.

**Extended Data Fig. 3:**
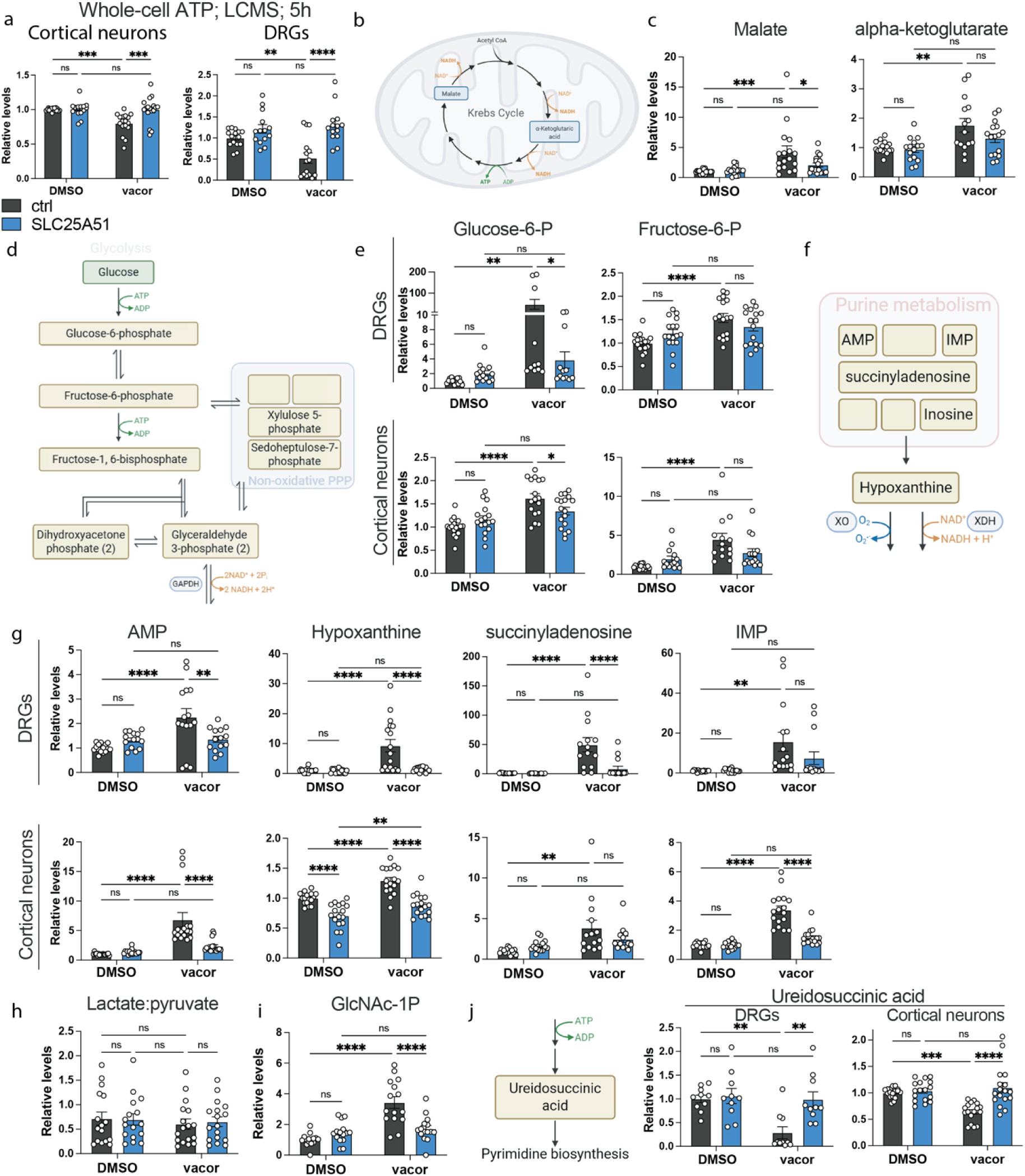
a. ATP levels in primary mouse dorsal root ganglia (DRG) and cortical neurons transduced with A51 vs control, 5 hours following treatment with vacor (12.5 µM for DRGs;2.5 µM for cortical neurons). Metabolite levels are shown relative to DMSO-treated control, as measured by LC-MS. Error bars show mean ± SEM, with data points representing independently treated culture wells (n=15-17) pooled across 3 independent culture preparations. Statistical significance was determined by 2-way ANOVA with Holm-Šídák’s multiple comparisons test. b. Schematic showing malate and alpha-ketoglutarate as products of NAD+-dependent reactions in the Krebs cycle in mitochondria. Created in BioRender. DiAntonio, A. (2026) https://BioRender.com/p4wjb9i. c. Malate and alpha-ketoglutarate levels in primary mouse dorsal root ganglia (DRG) and cortical neurons transduced with A51 vs control, 5 hours following treatment with vacor (12.5 µM for DRGs;2.5 µM for cortical neurons). Metabolite levels are shown relative to DMSO-treated control, as measured by LC-MS. Error bars show mean ± SEM, with data points representing independently treated culture wells (n=15-17) pooled across 3 independent culture preparations. Statistical significance was determined by 2-way ANOVA with Holm-Šídák’s multiple comparisons test. d. Schematic of the major steps in glycolysis, including the metabolite intermediates, the enzymes involved, and the ATP and NAD+-consuming steps. Created in BioRender. DiAntonio, A. (2026) https://BioRender.com/p4wjb9i. e. Levels of glycolytic intermediates glucose-6-phosphate and fructose-6-phosphate in DRGs and cortical neurons transduced with A51 vs control, 5 hours following treatment with vacor (12.5 µM for DRGs;2.5 µM for cortical neurons). Metabolite levels are shown relative to DMSO-treated control, as measured by LC-MS. Error bars show mean ± SEM, with data points representing independently treated culture wells (n=15 for DRGs; n=17 for cortical neurons) pooled across 3 independent culture preparations. Statistical significance was determined by 2-way ANOVA with Holm-Šídák’s multiple comparisons test. f. Schematic of purine intermediates upstream of NAD+ bottleneck. Created in BioRender. DiAntonio, A. (2026) https://BioRender.com/p4wjb9i. g. Levels of purine intermediates AMP, IMP, hypoxanthine, and succinyladenosine in DRGs and cortical neurons transduced with A51 vs control, 5 hours following treatment with vacor (12.5 µM for DRGs;2.5 µM for cortical neurons). Metabolite levels are shown relative to DMSO-treated control, as measured by LC-MS. Error bars show mean ± SEM, with data points representing independently treated culture wells (n=15 for DRGs; n=17 for cortical neurons) pooled across 3 independent culture preparations. Statistical significance was determined by 2-way ANOVA with Holm-Šídák’s multiple comparisons test. h. Lactate to pyruvate ratio in cortical neurons transduced with A51 vs control, 5 hours following treatment with vacor (2.5 µM). Metabolites were measured by LC-MS. Error bars show mean ± SEM, with data points representing independently treated culture wells (n=16-17) pooled across 3 independent culture preparations. Statistical significance was determined by 2-way ANOVA with Tukey’s multiple comparisons test. i. Hexosamine biosynthesis pathway intermediate N-acetylglucosamine 1-phosphate in DRGs transduced with A51 vs control, 5 hours following treatment with vacor (12.5 µM). Metabolite levels are shown relative to DMSO-treated control, as measured by LC-MS. Error bars show mean ± SEM, with data points representing independently treated culture wells (n=15) pooled across 3 independent culture preparations. Statistical significance was determined by 2-way ANOVA with Holm-Šídák’s multiple comparisons test. j. Schematic and levels of ureidosuccinic acid in primary mouse dorsal root ganglia (DRG) and cortical neurons transduced with A51 vs control, 5 hours following treatment with vacor (12.5 µM for DRGs;2.5 µM for cortical neurons). Metabolite levels are shown relative to DMSO-treated control, as measured by LC-MS. Error bars show mean ± SEM, with data points representing independently treated culture wells (n=10-17) pooled across 3 independent culture preparations. Statistical significance was determined by 2-way ANOVA with Holm-Šídák’s multiple comparisons test. Created in BioRender. DiAntonio, A. (2026) https://BioRender.com/p4wjb9i (left).

**Extended Data Fig. 4.**
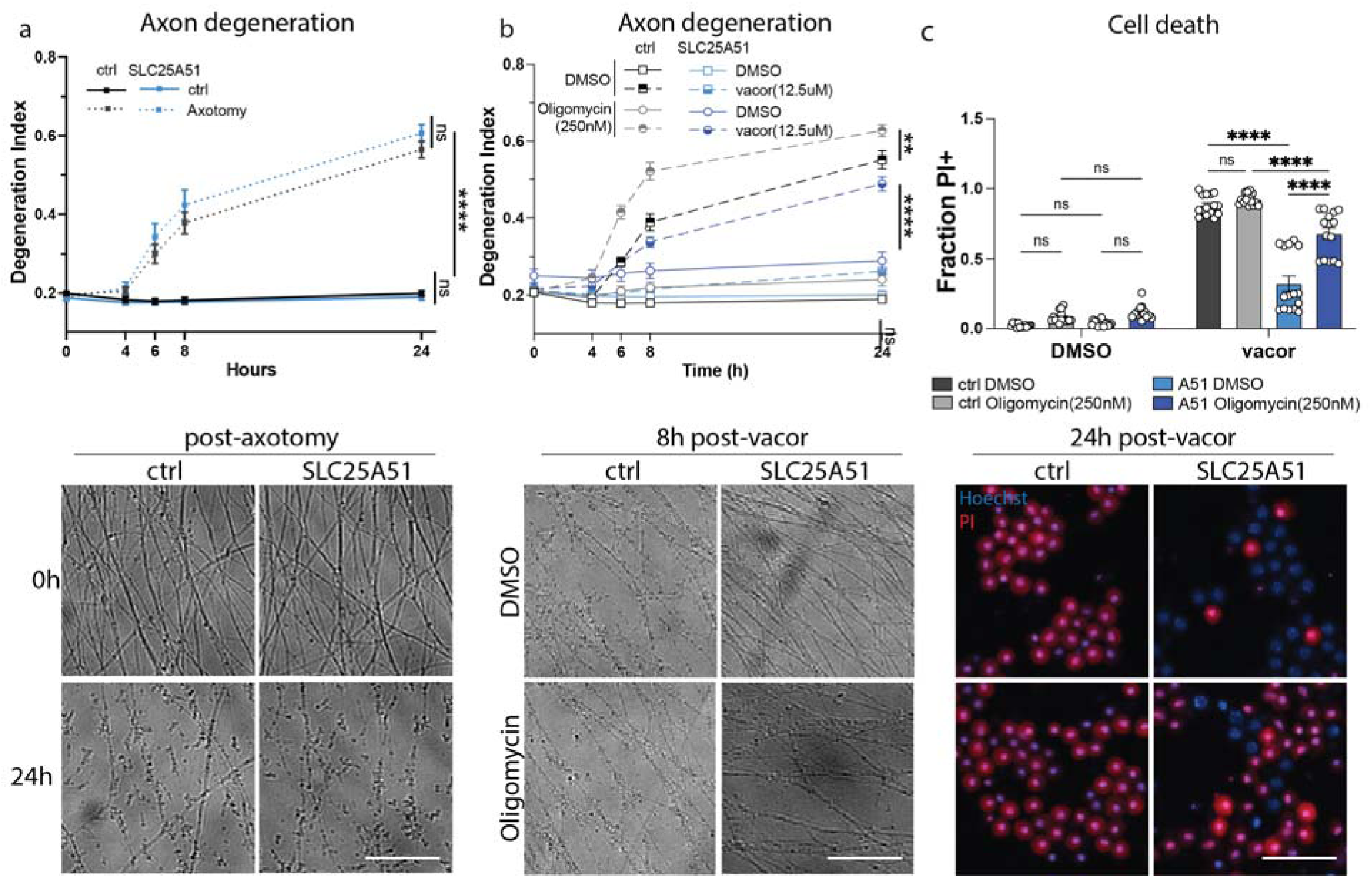
a. Axon degeneration in primary DRG neurons transduced with SLC25A51 (A51) vs control (empty vector or GFP) over 24 hours following axotomy. Degeneration index was quantified, with an increase indicating increased degeneration. Error bars show mean of independently treated culture wells, pooled across 3 independent culture preparations, ± SEM, with n=17-20. Statistical significance was determined by a 2-way ANOVA with Tukey’s multiple comparisons test with pairwise comparisons shown for 0/24h. Representative axon images are shown at 24 hours following axotomy in control (empty vector) vs SLC25A51-overexpressing neurons. Scale bar, 50 µm. b. Axon degeneration in primary DRG neurons transduced with A51 vs empty vector control, treated with vacor (12.5 µM) ± ATP synthase inhibitor oligomycin (250 nM) vs DMSO over 24 hours. Degeneration index was quantified, with an increase indicating increased degeneration. Error bars show mean of independently treated culture wells, pooled across 3 independent culture preparations, ± SEM, with n=15. Statistical significance was determined by 2-way ANOVA with Tukey’s multiple comparisons test with pairwise comparisons shown for 6h. Representative axon images are shown at 8 hours post-vacor in control (empty vector) vs SLC25A51-overexpressing neurons ± oligomycin. Scale bar, 50 µm. c. Cell death in primary DRG neurons transduced with A51 or control (GFP/empty vector) and treated with vacor (12.5 µM) ± ATP synthase inhibitor oligomycin (250 nM) vs DMSO for 24 hours, measured by the fraction of nuclei positive for propidium iodide (PI). Error bars show mean ± SEM, with data points representing independently treated culture wells (n=15) pooled across 3 independent culture preparations. Statistical significance was determined with a 2-way ANOVA with Tukey’s multiple comparisons test. Representative images showing Hoechst+ nuclei (blue) and propidium iodide (PI) positive dead cells (red) at 24 hours post vacor. Scale bar, 50 µm.

**Extended Data Fig. 5.**
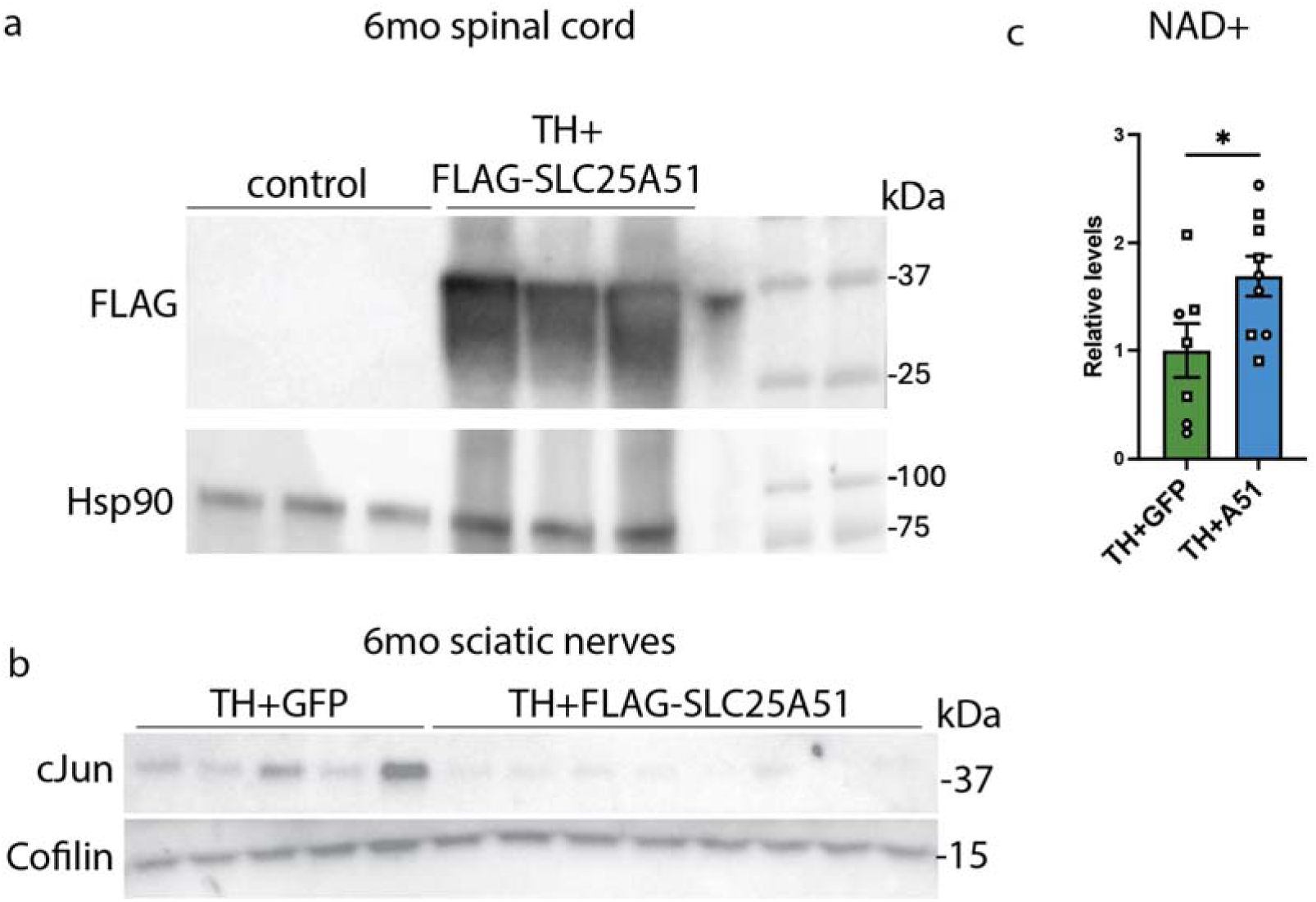
a. Western blot showing the presence of FLAG in 6mo spinal cord lysates from TH mice injected with AAV-FLAG-SLC25A51 but not in uninjected TH controls, with Hsp90 as loading control. b. Full western blot confirmation of reduced levels of repair Schwann cell marker c-Jun in 6-month sciatic nerves of TH mice injected with AAV-FLAG-SLC25A51 vs AAV-GFP control. c. NAD+ levels in sciatic nerves of TH mice injected with AAV-SLC25A51 vs AAV-GFP, measured at 6 months of age. Metabolite levels are shown relative to GFP-treated control, as measured by LC-MS. Error bars show mean ± SEM, with data points representing individual animals (n=7 for GFP; n=10 for A51). Statistical significance was determined with Welch’s t-test. Males are represented as squares and females as circles.

## References

1. Essuman, K. et al. The SARM1 Toll/Interleukin-1 Receptor Domain Possesses Intrinsic NAD+ Cleavage Activity that Promotes Pathological Axonal Degeneration. Neuron 93, 1334–1343.e5 (2017).

2. Osterloh, J. M. et al. dSarm/Sarm1 Is Required for Activation of an Injury-Induced Axon Death Pathway. Science 337, 481–484 (2012).

3. Gerdts, J., Summers, D. W., Sasaki, Y., DiAntonio, A. & Milbrandt, J. Sarm1-mediated axon degeneration requires both SAM and TIR interactions. Journal of Neuroscience 33, 13569–13580 (2013).

4. Wu, T. et al. SARM1 executes neuronal parthanatos and promotes excitotoxic cell death. Neuron 114, 2975–2986.e5 (2026).

5. Ding, C. et al. SARM1 loss protects retinal ganglion cells in a mouse model of autosomal dominant optic atrophy. J Clin Invest 135, (2025).

6. Sato-Yamada, Y. et al. A SARM1-mitochondrial feedback loop drives neuropathogenesis in a Charcot-Marie-Tooth disease type 2A rat model. J Clin Invest 132, e161566 (2022).

7. Henninger, N. et al. Attenuated traumatic axonal injury and improved functional outcome after traumatic brain injury in mice lacking Sarm1. Brain 139, 1094–1105 (2016).

8. Geisler, S. et al. Prevention of vincristine-induced peripheral neuropathy by genetic deletion of SARM1 in mice. Brain 139, 3092–3108 (2016).

9. White, M. A. et al. Sarm1 deletion suppresses TDP-43-linked motor neuron degeneration and cortical spine loss. Acta Neuropathol Commun 7, 166 (2019).

10. Miao, X. et al. SARM1 Promotes Neurodegeneration and Memory Impairment in Mouse Models of Alzheimer’s Disease. Aging and disease 15, 390–407 (2024).

11. Jiang, Y. et al. The NAD+-mediated self-inhibition mechanism of pro-neurodegenerative SARM1. Nature 588, 658–663 (2020).

12. Figley, M. D. et al. SARM1 is a metabolic sensor activated by an increased NMN/NAD+ ratio to trigger axon degeneration. Neuron 109, 1118–1136.e11 (2021).

13. Gerdts, J., Brace, E. J., Sasaki, Y., DiAntonio, A. & Milbrandt, J. SARM1 activation triggers axon degeneration locally via NAD+ destruction. Science 348, 453–457 (2015).

14. Angeletti, C. et al. SARM1 is a multi-functional NAD(P)ase with prominent base exchange activity, all regulated bymultiple physiologically relevant NAD metabolites. iScience 25, 103812 (2022).

15. Hinz, F. I. & Hoogenraad, C. C. DLK, NMNAT2, and SARM1: Judge, Jury, and Executioner in Axon Degeneration. Annu Rev Biochem 95, 595–624 (2026).

16. Loreto, A. & Neukomm, L. J. Programmed axon degeneration: mechanism, inhibition and therapeutic potential. Nat. Rev. Neurosci. 27, 44–60 (2026).

17. Coleman, M. P. & Höke, A. Programmed axon degeneration: from mouse to mechanism to medicine. Nat Rev Neurosci 21, 183–196 (2020).

18. Pan, W., Guo, D., Liu, D. & Wang, X. A metabolic cell death program downstream of SARM1 couples NAD+ depletion to BAX activation and APAF1 degradation. Proc Natl Acad Sci U S A 122, e2522444122 (2025).

19. Shi, Y. et al. Structural basis of SARM1 activation, substrate recognition, and inhibition by small molecules. Molecular Cell 10.1016/j.molcel.2022.03.007 (2022) doi:10.1016/j.molcel.2022.03.007.

20. Bratkowski, M. et al. Uncompetitive, adduct-forming SARM1 inhibitors are neuroprotective in preclinical models of nerve injury and disease. Neuron 110, 3711–3726.e16 (2022).

21. Hughes, R. O. et al. Small Molecule SARM1 Inhibitors Recapitulate the SARM1^-/-^ Phenotype and Allow Recovery of a Metastable Pool of Axons Fated to Degenerate. Cell Rep 34, 108588 (2021).

22. Alano, C. C. et al. Differences among cell types in NAD(+) compartmentalization: a comparison of neurons, astrocytes, and cardiac myocytes. J Neurosci Res 85, 3378–3385 (2007).

23. Cambronne, X. A. et al. NAD+ biosensor reveals multiple sources for mitochondrial NAD+. Science 352, 1474–1477 (2016).

24. Cambronne, X. A. & Kraus, W. L. Location, Location, Location: Compartmentalization of NAD+ Synthesis and Functions in Mammalian Cells. Trends Biochem Sci 45, 858–873 (2020).

25. Høyland, L. E. et al. Subcellular NAD+ pools are interconnected and buffered by mitochondrial NAD+. Nat Metab 10.1038/s42255-024-01174-w (2024) doi:10.1038/s42255-024-01174-w.

26. Kory, N. et al. MCART1/SLC25A51 is required for mitochondrial NAD transport. Science Advances 6, eabe5310 (2020).

27. Luongo, T. S. et al. SLC25A51 is a mammalian mitochondrial NAD+ transporter. Nature 588, 174–179 (2020).

28. Girardi, E. et al. Epistasis-driven identification of SLC25A51 as a regulator of human mitochondrial NAD import. Nat Commun 11, 6145 (2020).

29. Nikiforov, A., Dölle, C., Niere, M. & Ziegler, M. Pathways and Subcellular Compartmentation of NAD Biosynthesis in Human Cells. J Biol Chem 286, 21767– 21778 (2011).

30. Berger, F., Lau, C., Dahlmann, M. & Ziegler, M. Subcellular compartmentation and differential catalytic properties of the three human nicotinamide mononucleotide adenylyltransferase isoforms. J Biol Chem 280, 36334–36341 (2005).

31. VanLinden, M. R. et al. Subcellular Distribution of NAD+ between Cytosol and Mitochondria Determines the Metabolic Profile of Human Cells. J Biol Chem 290, 27644–27659 (2015).

32. Loreto, A. et al. Neurotoxin-mediated potent activation of the axon degeneration regulator SARM1. eLife 10, e72823 (2021).

33. Sasaki, Y. et al. cADPR is a gene dosage-sensitive biomarker of SARM1 activity in healthy, compromised, and degenerating axons. Experimental Neurology 329, (2020).

34. Zou, Y. et al. Illuminating NAD+ Metabolism in Live Cells and In Vivo Using a Genetically Encoded Fluorescent Sensor. Dev Cell 53, 240–252.e7 (2020).

35. Ko, K. W., Devault, L., Sasaki, Y., Milbrandt, J. & DiAntonio, A. Live imaging reveals the cellular events downstream of SARM1 activation. eLife 10, e71148 (2021).

36. Marvin, J. S. et al. iATPSnFR2: A high-dynamic-range fluorescent sensor for monitoring intracellular ATP. Proc Natl Acad Sci U S A 121, e2314604121 (2024).

37. Baur, J., Luongo, T., Eller, J., Lu, M. & Cambronne, X. SLC25A51 is a Mitochondrial NAD Transporter. The FASEB Journal 35, (2021).

38. Todisco, S., Agrimi, G., Castegna, A. & Palmieri, F. Identification of the Mitochondrial NAD+ Transporter in Saccharomyces cerevisiae. Journal of Biological Chemistry 281, 1524–1531 (2006).

39. Sasaki, Y., Nakagawa, T., Mao, X., DiAntonio, A. & Milbrandt, J. NMNAT1 inhibits axon degeneration via blockade of SARM1-mediated NAD+ depletion. eLife 5, e19749 (2016).

40. Yamamoto, M. et al. Nmnat3 Is Dispensable in Mitochondrial NAD Level Maintenance In Vivo. PLoS ONE 11, e0147037 (2016).

41. Hikosaka, K. et al. Deficiency of Nicotinamide Mononucleotide Adenylyltransferase 3 (Nmnat3) Causes Hemolytic Anemia by Altering the Glycolytic Flow in Mature Erythrocytes*. Journal of Biological Chemistry 289, 14796–14811 (2014).

42. Gilley, J. & Coleman, M. P. Endogenous Nmnat2 Is an Essential Survival Factor for Maintenance of Healthy Axons. PLoS Biology 8, (2010).

43. Gilley, J., Orsomando, G., Nascimento-Ferreira, I. & Coleman, M. P. Absence of SARM1 rescues development and survival of NMNAT2-Deficient axons. Cell Reports 10, 1975–1982 (2015).

44. Dingwall, C. B. et al. Macrophage depletion blocks congenital SARM1-dependent neuropathy. J Clin Invest 132, e159800 (2022).

45. Summers, D. W., DiAntonio, A. & Milbrandt, J. Mitochondrial dysfunction induces Sarm1-dependent cell death in sensory neurons. J Neurosci 34, 9338–9350 (2014).

46. Loreto, A. et al. Mitochondrial impairment activates the Wallerian pathway through depletion of NMNAT2 leading to SARM1-dependent axon degeneration. Neurobiology of Disease 134, 104678 (2020).

47. Loreto, A. et al. SARM1 activation induces reversible mitochondrial dysfunction and can be prevented in human neurons by antisense oligonucleotides. Neurobiol Dis 213, 106986 (2025).

48. Figley, M. D. & DiAntonio, A. The SARM1 axon degeneration pathway: control of the NAD+ metabolome regulates axon survival in health and disease. Current Opinion in Neurobiology 63, 59–66 (2020).

49. Zha, M. et al. Molecular mechanism of ADP-ribose hydrolysis by human NUDT5 from structural and kinetic studies. J Mol Biol 379, 568–578 (2008).

50. Perraud, A.-L. et al. NUDT9, a member of the Nudix hydrolase family, is an evolutionarily conserved mitochondrial ADP-ribose pyrophosphatase. J Biol Chem 278, 1794–1801 (2003).

51. van Heerden, J. H. et al. Lost in Transition: Start-Up of Glycolysis Yields Subpopulations of Nongrowing Cells. Science 343, 1245114 (2014).

52. Bell, S. et al. Lesch-Nyhan disease causes impaired energy metabolism and reduced developmental potential in midbrain dopaminergic cells. Stem Cell Reports 16, 1749–1762 (2021).

53. Avetisyan, A. et al. A dNmnat-sensitized in vivo platform for unbiased discovery of regulators of neurodegeneration. 2025.09.25.678663 Preprint at 10.1101/2025.09.25.678663 (2025).

54. Ali, Y. O., Li-Kroeger, D., Bellen, H. J., Zhai, R. G. & Lu, H.-C. NMNATs, Evolutionarily Conserved Neuronal Maintenance Factors. Trends Neurosci 36, 10.1016/j.tins.2013.07.002 (2013).

55. Ratia, K. M. et al. A mechanism of allosteric modulation of nicotinamide phosphoribosyltransferase to elevate cellular NAD+. Biochemistry 62, 923–933 (2023).

56. Mukherjee, S. et al. Hepatocyte mitochondrial NAD+ content is limiting for liver regeneration. Nat Metab 1–14 (2025) doi:10.1038/s42255-025-01408-5.

57. Wang, X. et al. Mitochondrial NAD+ Transport Alleviates Cerebral Ischemia/Reperfusion Injury via Enhancement of Mitochondrial Function. Antioxidants & Redox Signaling 45, 359–375 (2026).

58. Yang, H. et al. Nutrient-Sensitive Mitochondrial NAD+ Levels Dictate Cell Survival. Cell 130, 1095–1107 (2007).

59. Goyal, S., Lyons, S. N. & Cambronne, X. A. Mitochondrial NAD^+^ gradient sustained by membrane potential and transport. Sci. Adv. 11, eaea7460 (2025).

60. Sasaki, Y. et al. SARM1 depletion rescues NMNAT1-dependent photoreceptor cell death and retinal degeneration. eLife 9, e62027 (2020).

61. Lois, C., Hong, E. J., Pease, S., Brown, E. J. & Baltimore, D. Germline transmission and tissue-specific expression of transgenes delivered by lentiviral vectors. Science 295, 868–872 (2002).

62. Sasaki, Y., Vohra, B. P. S., Lund, F. E. & Milbrandt, J. Nicotinamide Mononucleotide Adenylyl Transferase-Mediated Axonal Protection Requires Enzymatic Activity But Not Increased Levels of Neuronal Nicotinamide Adenine Dinucleotide. J. Neurosci. 29, 5525–5535 (2009).

63. Sasaki, Y., Nakagawa, T., Mao, X., Diantonio, A. & Milbrandt, J. NMNAT1 inhibits axon degeneration via blockade of SARM1-mediated NAD + depletion. 10.7554/eLife.19749.001 (2016) doi:10.7554/eLife.19749.001.

64. Sasaki, Y., Hackett, A. R., Kim, S., Strickland, A. & Milbrandt, J. Dysregulation of NAD+ Metabolism Induces a Schwann Cell Dedifferentiation Program. J Neurosci 38, 6546–6562 (2018).

65. Pang, Z. et al. MetaboAnalyst 6.0: towards a unified platform for metabolomics data processing, analysis and interpretation. Nucleic Acids Res 52, W398–W406 (2024).

66. Krus, K. L. et al. Two cardinal features of ALS, reduced STMN2 and pathogenic TDP-43, synergize to accelerate motor decline in mice. Exp Neurol 384, 115068 (2025).

67. Kaiser, T. et al. MyelTracer: A Semi-Automated Software for Myelin g-Ratio Quantification. eNeuro 8, (2021).

